# Regulatory Network Inference of Induced Senescent Midbrain Cell Types Reveals Cell Type-Specific Senescence-Associated Transcriptional Regulators

**DOI:** 10.1101/2025.02.06.636893

**Authors:** Taylor Russo, Jonathan Plessis-Belair, Roger Sher, Markus Riessland

**Affiliations:** Department of Neurobiology and Behavior; Stony Brook University, Stony Brook, NY 11794, USA; Center for Nervous System Disorders; Stony Brook University, Stony Brook, NY 11794, USA

## Abstract

Cellular senescence of brain cell types has become an increasingly important perspective for both aging and neurodegeneration, specifically in the context of Parkinson’s Disease (PD). The characterization of classical hallmarks of senescence is a widely debated topic, whereby the context in which a senescence phenotype is being investigated, such as the cell type, the inducing stressor, and/or the model system, is an extremely important aspect to consider when defining a senescent cell. Here, we describe a cell type-specific profile of senescence through the investigation of various canonical senescence markers in five human midbrain cell lines using chronic 5-Bromodeoxyuridine (BrdU) treatment as a model of DNA damage-induced senescence. We used principal component analysis (PCA) and subsequent regulatory network inference to define both unique and common senescence profiles in the cell types investigated, as well as revealed senescence-associated transcriptional regulators (SATRs). Functional characterization of one of the identified regulators, transcription factor AP4 (TFAP4), further highlights the cell type-specificity of the expression of the various senescence hallmarks. Our data indicates that SATRs modulate cell type-specific profiles of induced senescence in key midbrain cell types that play an important role in the context of aging and PD.

## Introduction

Parkinson’s Disease (PD) is the second most common age-related neurodegenerative disorder and is characterized by the progressive loss of dopaminergic (DA) neurons in the substantia nigra pars compacta (SNpc) (*1*). Although age is the biggest risk factor for PD and the degeneration of DA neurons is apparent, the exact cause(s) of their loss is unclear and no current treatment options exist to prevent or slow the DA neuron degeneration or the development of related symptoms. An emerging view, however, is that cellular senescence pathways are key mechanisms contributing to DA neuron loss in PD (*2*). Cellular senescence is a state of irreversible cell-cycle arrest in mitotic cells resulting from long-term exposure to DNA damage or oxidative stress (*3*). Senescence is characterized by a variety of key hallmarks, including proliferation arrest, increased senescence-associated β-galactosidase (β-gal) activity, elevated levels of cyclin-dependent kinase (CDK) inhibitors p21/p16/p19, expression of the senescence-associated secretory phenotype (SASP), senescence-associated heterochromatin foci (SAHF), mitochondrial and/or lysosomal dysfunction, and morphological changes including increased nuclear size due to loss of nuclear envelope protein lamin B1 (*4–7*). Importantly, not all of these phenotypes are present in every senescent cell or model of senescence, and there are widely reported differences in the expression of these features based on the method of senescence induction, the cell type being studied, and the timepoint at which senescence is assayed (*8*).

Recent advancements have been made to provide a ‘minimum criteria’ to define a senescent cell *in vivo* based on observed phenotypes (*8*), however senescent cells *in vivo* are very low in number and transient in their presence as they can be cleared within days by immune cells (*9, 10*). In contrast, *in vitro* models of senescence are advantageous where an unlimited number of cells can be generated and molecular investigations into the nuanced differences in the expression of senescent phenotypes across different cell types can be made more accessible. Additionally, recent investigations have determined that senescence occurs in post-mitotic cells of the brain, such as cortical neurons in Alzheimer’s Disease (AD) and DA neurons in human and mouse models of PD, and can spread to other brain cell types in both normal aging and in the context of neurodegenerative diseases (*11–13*). Thus, it is plausible that a spreading of senescence between diverse brain cell types could underlie various neurodegenerative phenotypes (*4*).

The characterization of cell type-specific senescence would allow for more precise, effective, and safe therapeutic interventions, particularly in complex diseases like PD where evidence suggests that multiple cell types contribute to pathology (*14*). Additionally, this would also help to unravel the dual roles of senescence in health and disease, ensuring that therapies target the harmful nature of senescent cells while preserving their more beneficial functions. Specifically, a better understanding of the senescence profiles of various midbrain cell types *in vitro* would bolster future *in vivo* investigations of senescence in the context of aging and PD. We sought to systematically characterize an established method of senescence induction in human cell lines to delineate a cell type-specific profile of senescence. We utilized chronic 5-Bromodeoxyuridine (BrdU) treatment, which is an established inducer of classical replicative senescence hallmarks in various cell types (*15, 16*). This treatment was performed in five human cell lines to model relevant cell types of the midbrain, including astrocytes, endothelial cells, microglia, oligodendrocytes, and DA-like neurons. Importantly, our findings highlight cell type-specific profiles of key hallmarks displayed in cellular senescence. Additionally, the transcriptional profile of the different senescent cell lines was sufficient to isolate a unifying senescence profile independent of cell identity. We then utilized regulatory network inference (*17*) to reveal transcriptional regulators of this senescence phenotype across the midbrain cell types (SATRs) and observed that knockdown of select regulators induced varying senescence hallmarks based on the cell type being investigated. We then specifically focused on TFAP4 because of its relevant biological function, where it has been previously shown to suppress senescence in colorectal cancer cells (*18*). We performed siRNA knockdowns (KD) in the five midbrain cell types and demonstrated that based on the cell type being investigated, there is a specific expression of the different senescence hallmarks when TFAP4 is repressed. Taken together, we have established that chronic BrdU treatment of human midbrain immortalized cell lines is a reliable and reproducible *in vitro* model of senescence. Additionally, we have isolated a senescence transcriptional signature that shares common upstream transcriptional regulators (SATRs), which themselves modulate specific hallmarks of senescence dependent on the cell type.

## Results

### Induction of a canonical senescence phenotype in human midbrain cell lines

To identify a senescence profile in each of the midbrain cell types, we used a common DNA damage-inducing stressor to characterize hallmarks of senescence. We treated with 100 uM BrdU (*19*) or a DMSO control for 7 days in SVG-A (astrocytes), HBEC-5i (endothelial cells), HMC3 (microglia), HOG (oligodendrocytes), and SK-N-MC (neurons with some DA-like features, such as moderate dopamine-beta-hydroxylase activity) cell lines and analyzed various well-established and widely accepted hallmarks of a senescence phenotype (Figure 1A). First, we observed a significant increase in the percentage of senescence-associated β-gal positive cells following 7-day BrdU treatment across all five midbrain cell types (Figure 1B). Next, we characterized the proliferation arrest of BrdU-treated cells, a defining hallmark of senescence. We measured the growth rates (divisions/hour) across treatment groups for 48 hours and observed a halt in proliferation following BrdU treatment in all cell types (Figure 1C, S1A). We went on to measure γH2AX (pS139) foci (senescence-associated heterochromatin foci; SAHF) as a molecular senescence-associated marker of DNA damage. We observed a significant increase in the number of γH2AX foci per cell following BrdU treatment in astrocytes, endothelial cells, and neurons (Figure 1D, S1B). Next, we characterized nuclear size using staining for lamin B1 and observed a significant increase in nuclear area with BrdU treatment across all cell types except microglia (Figure 1E, 2C). Interestingly, staining for CDK inhibitor and canonical early-stage senescence marker p21 (*6*) in BrdU-treated cell types showed a specific increase in astrocytes, endothelial cells, and oligodendrocytes, but no significant p21 upregulation was observed in microglia or in neurons (Figure 1F, S1B).

**Figure 1:**
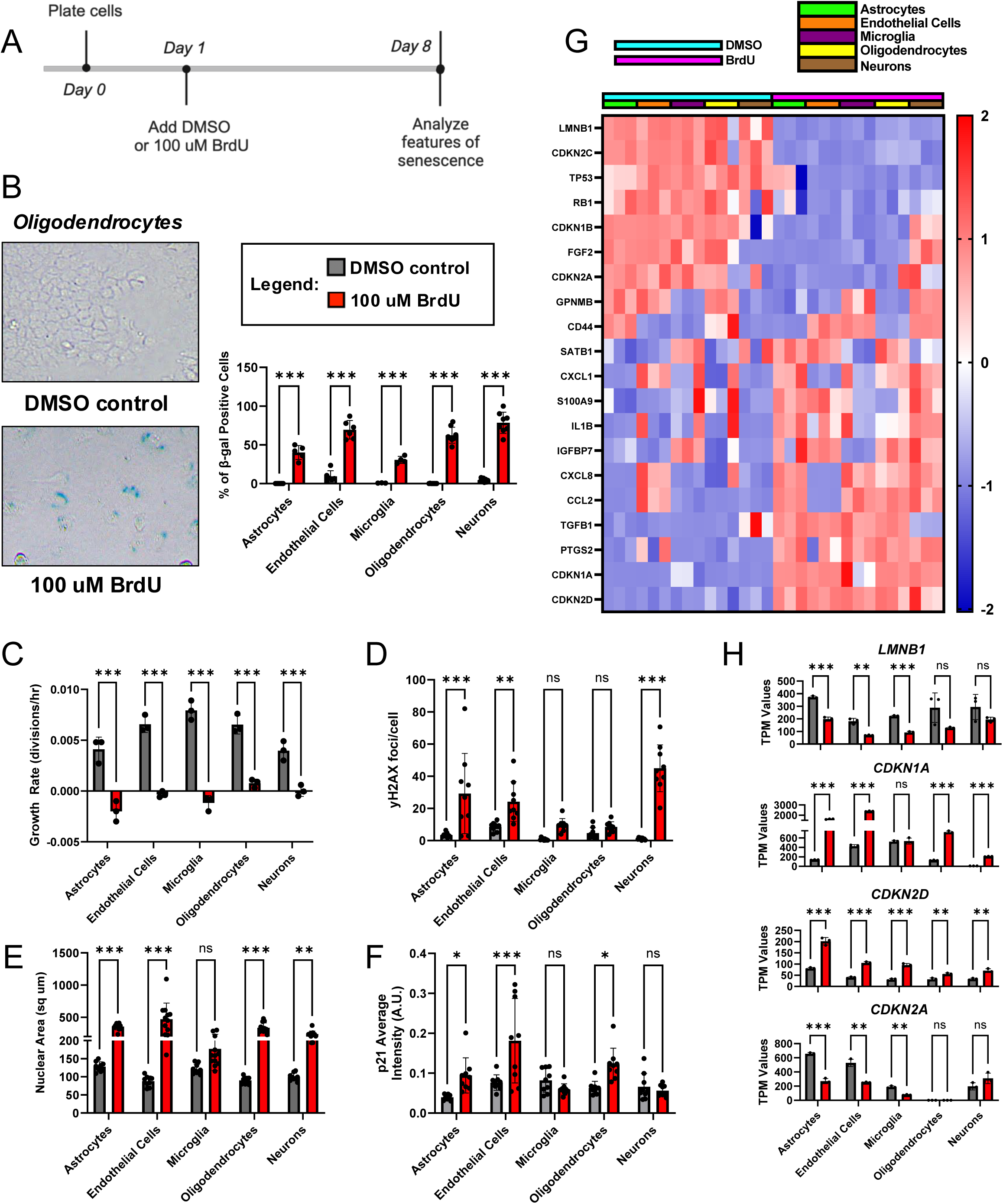
Induction of a canonical senescence phenotype in human midbrain cell lines. A) Timeline of treatment of human cell lines with DMSO control or 100 uM BrdU for 7 days. Features of senescence were analyzed 8 days total after initial plating of cells. B) Representative images of senescence-associated ß-galactosidase (β-gal) staining in oligodendrocytes after 7-day treatment with either DMSO control or 100 uM BrdU. Quantification of percentage of β-gal positive cells (blue) in astrocytes, endothelial cells, microglia, oligodendrocytes, and neurons following 7-day treatment with DMSO control (grey) or 100 uM BrdU (red) (n=3-8). C) Growth rates (divisions/hour) of DMSO (grey) and 100 uM BrdU (red) treated human cell lines (n=3). D) Quantification of immunofluorescence staining of DNA damage marker γH2AX in DMSO (grey) and 100 uM BrdU (red) treated human cell lines (n=9). E) Quantification of nuclear area based on immunofluorescence staining of lamin B1 in DMSO (grey) and 100 uM BrdU (red) treated human cell lines (n=9). F) Quantification of immunofluorescence staining of canonical senescence marker p21 in DMSO (grey) and 100 uM BrdU (red) treated human cell lines (n=9). G) Z-score heatmap showing canonical senescence marker expression in all cell types (astrocytes – green, endothelial cells – orange, microglia – purple, oligodendrocytes – yellow, neurons – brown) based on bulk RNAseq of DMSO (blue) and BrdU (pink) treated human cell lines. H) TPM expression values of highlighted senescence markers *LMNB1*, *CDKN1A*, *CDKN2D*, and *CDKN2A* in DMSO (grey) and BrdU (red) treated cell lines. (n=3). Data was analyzed by two-way ANOVA with Šídák’s multiple comparisons test. All graphs show mean with error bars depicting standard deviation (ns, p>0.05, * p<0.05, ** p<0.01, *** p<0.001).

**Figure 2:**
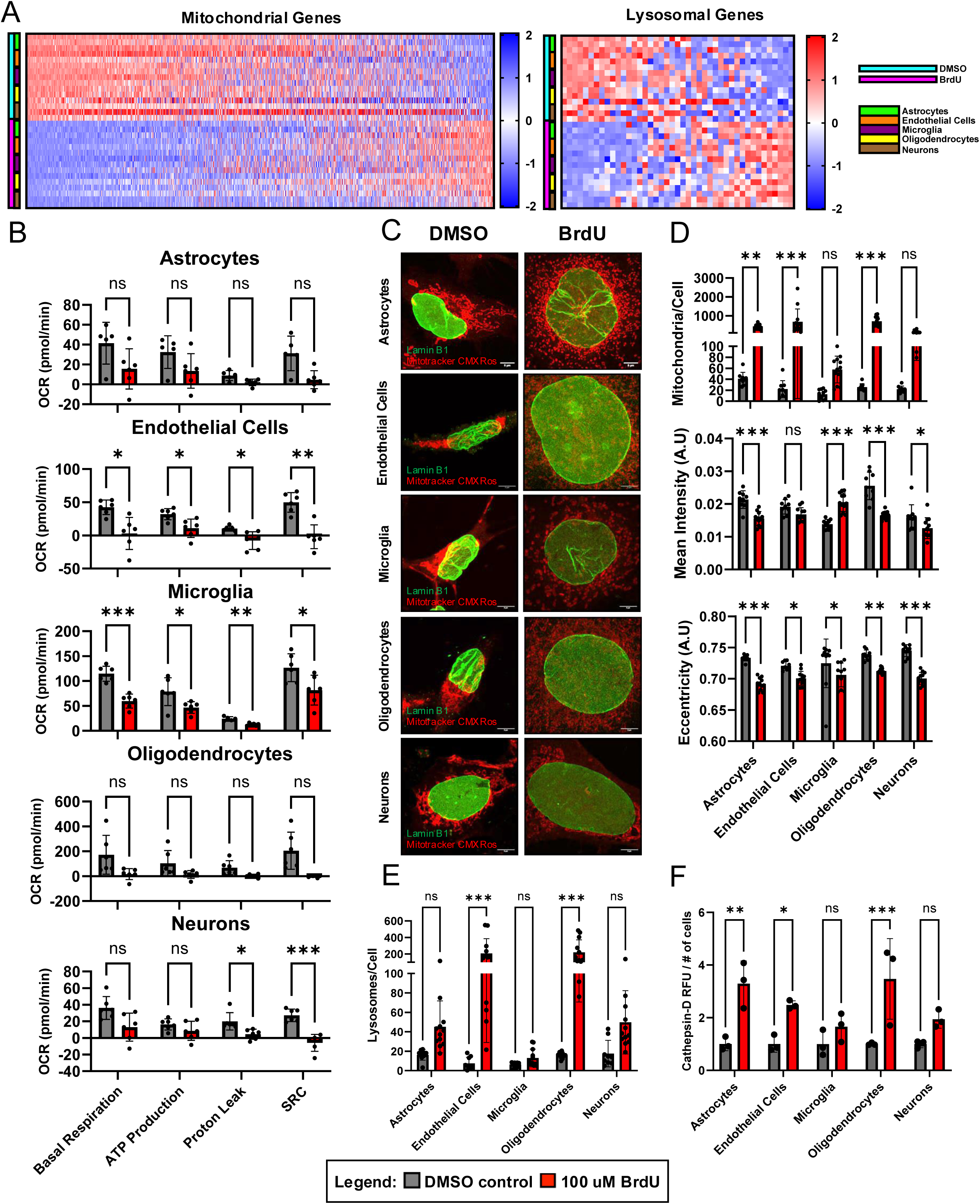
Induced senescent human midbrain cell lines show mitochondrial and lysosomal alterations. A) Z-score heatmaps showing altered expression of mitochondrial and lysosomal genes following BrdU treatment (pink) as compared to DMSO control (blue) in human cell lines (astrocytes – green, endothelial cells – orange, microglia – purple, oligodendrocytes – yellow, neurons – brown). B) Quantification of alterations in oxygen consumption rate from Seahorse XF Mito Stress test following 7-day treatment with DMSO (grey) or 100 uM BrdU (red) showing ATP production, proton leak, basal respiration, and spare respiratory capacity (SRC) across astrocytes, endothelial cells, microglia, oligodendrocytes, and neurons (top-bottom, n=5-6). C) Airyscan confocal images of human cell lines following DMSO and BrdU treatment with staining for lamin B1 (green) and Mitotracker Red CMXRos (red). D) Quantification of Mitotracker Red CMX Ros staining including mitochondrial mass (top), eccentricity (indicative of increased mitochondrial fission) (middle), and mitochondrial membrane potential (based on intensity of Mitotracker staining) (bottom) from 7-day DMSO control (grey) and 100 uM BrdU treated (red) (n=9-12). E) Quantification of Lysotracker Deep Red staining (n=9-12). F) Cathepsin D activity following 100 uM BrdU treatment (red) in all cell types as compared to DMSO control (grey) (n=3). Data from (B) analyzed by multiple unpaired two-tailed t-tests. Data from (D-F) analyzed by two-way ANOVA with Šídák’s multiple comparisons test. All graphs show mean with standard deviation (ns, p>0.05, * p<0.05, ** p<0.01, *** p<0.001). Scale bars are 5 um.

To further characterize our BrdU-induced model of senescence, we performed bulk RNAseq of DMSO and BrdU-treated samples from all five cell lines. Extracting canonical senescence differentially expressed genes (DEGs), we revealed both common and unique transcriptional alterations following senescence induction (Figure 1G). Specifically, nuclear envelope marker *LMNB1* was decreased across all cell types with BrdU treatment, but only to significant levels in astrocytes, endothelial cells, and microglia (Figure 1H). Focusing on genes encoding CDK inhibitors, we observed that *CDKN1A* (p21) was significantly increased in all cell types except microglia with BrdU treatment (Figure 1H). Alternatively, microglia – along with all other cell types – demonstrated a significant increase in the senescence driver *CDKN2D* (p19) (*20*). Surprisingly, *CDKN2A* (p16) was significantly downregulated in astrocytes, endothelial cells, and microglia, but was unchanged in neurons and is not expressed in oligodendrocytes (Figure 1H). These transcriptional alterations indicate the induction of an early-stage senescence phenotype characterized by p21 and/or p19 expression rather than an upregulation of p16, which contributes to longer term maintenance of senescence (*6*). We also analyzed SASP marker expression in DMSO and BrdU-treated cells (Figure S1C) which showed a wide range of upregulated markers, as well as cell type differences in up- or down-regulation of various SASP factors. We went on to further validate p21 expression levels via Western blotting and detected significant increases in BrdU treated astrocytes, endothelial cells, and oligodendrocytes but not in microglia, consistent with observed transcriptional alterations. Neurons show upregulated yet insignificant p21 protein expression with BrdU treatment (Figure S1D). Taken together, these results demonstrate that BrdU-induced DNA damage was sufficient to induce senescence-associated β-gal activity and proliferation arrest in all five cell types. Investigations into nuclear size and canonical senescence-associated differentially expressed genes revealed cell type-specific differences, which were further elucidated through investigations into CDK inhibitor p21.

### Induced senescent human midbrain cell lines show mitochondrial and lysosomal alterations

We continued our investigation into key senescence hallmarks through the characterization of mitochondrial and lysosomal dysfunction. Analysis of mitochondrial and lysosomal gene lists based on our RNAseq dataset (Figure 2A) demonstrated a widespread transcriptional dysregulation of both mitochondrial and lysosomal genes with BrdU treatment (*21, 22*). We then performed functional assays to unravel the mitochondrial changes in each senescent cell type using measurements of the oxygen consumption rate (OCR) based on a Seahorse XF Analyzer and Mito Stress Test (Figure S2A) in DMSO and BrdU-treated cells (Figure 2B). Basal respiration and ATP production were significantly decreased only in senescent endothelial cells and microglia. Proton leak and spare respiratory capacity (SRC) were significantly decreased only in senescent microglia and neurons (Figure 2B). Interestingly, senescent astrocytes and oligodendrocytes showed overall diminished OCR, although the individual parameters were insignificant. Using MitoTracker Red CMXRos staining (Figure 2C-D, S2B), we observed a significant increase in mitochondrial mass (mitochondria/cell) in senescent astrocytes, endothelial cells, and oligodendrocytes (Figure 2D). Quantification of the mean intensity of MitoTracker Red CMXRos revealed a significant decrease in senescent astrocytes, oligodendrocytes, and neurons, highlighting a decrease in mitochondrial membrane potential. However, senescent microglia showed a significant increase in mitochondrial membrane potential based on MitoTracker Red CMXRos staining intensity. The mitochondria in the senescent cells also displayed a significantly reduced eccentricity based on their punctate appearance, as compared to the tubular, elongated mitochondria seen in the control cells (Figure 2D). Overall, these changes in mitochondrial morphology and dynamics show both similarities and differences across the senescent cell types.

Lastly, we functionally characterized changes in lysosomal content and function in the five cell lines following senescence induction. We used LysoTracker Deep Red staining to quantify the number of lysosomes per cell and saw a significant increase in senescent endothelial cells and oligodendrocytes, indicating a lack of lysosomal turnover (Figure 2E, S2C), which is in line with a characteristic increase in lysosomal mass as seen in other senescent cell models (*23*). Additionally, we used a cathepsin D activity assay to monitor lysosomal function and saw a significant increase in cathepsin D activity in senescent astrocytes, endothelial cells, and oligodendrocytes, with a trending increase in microglia and neurons (Figure 2F). Considering this characterization of mitochondrial and lysosomal alterations as well as the previously described hallmarks of senescence in Figure 1, we further highlight the ability of BrdU-induced DNA damage to produce both common and cell type-specific senescence phenotypes in midbrain cell types.

### Induced senescent human midbrain cell lines show cell type-specific responses to senolytic treatment

As a final investigation of the similarities and differences amongst the cell types in the context of DNA damage-induced senescence, we characterized the responsiveness of each cell type to well established senolytic drugs: dasatinib (D) + quercetin (Q) and navitoclax (ABT-263) (*24, 25*). We tested viability of all five cell types by exposing senescent cells to either 50 nM or 100 nM D + 10 uM or 20 uM Q and 1 uM, 10 uM, or 20 uM navitoclax for 48 hours. We observed that senescent oligodendrocytes were the most sensitive to D+Q (Figure 3D), whereas senescent astrocytes and microglia required a higher D+Q concentration to be effectively targeted (Figure 3A, C). Interestingly, senescent neurons and endothelial cells were relatively resistant to the same concentrations of D+Q (Figure 3B, E). Senescent astrocytes and oligodendrocytes were the most sensitive to the tested concentrations of navitoclax (Figure 3F, I), followed by microglia (Figure 3H). Navitoclax treatment in senescent endothelial cells and neurons was generally less effective (Figure 3G, J), where loss of non-senescent cells began at the same drug concentrations. Overall, these senolytic treatments emphasize the cell type-specificity in targeting senescent cells with senolytics, across the five different midbrain cell types.

**Figure 3:**
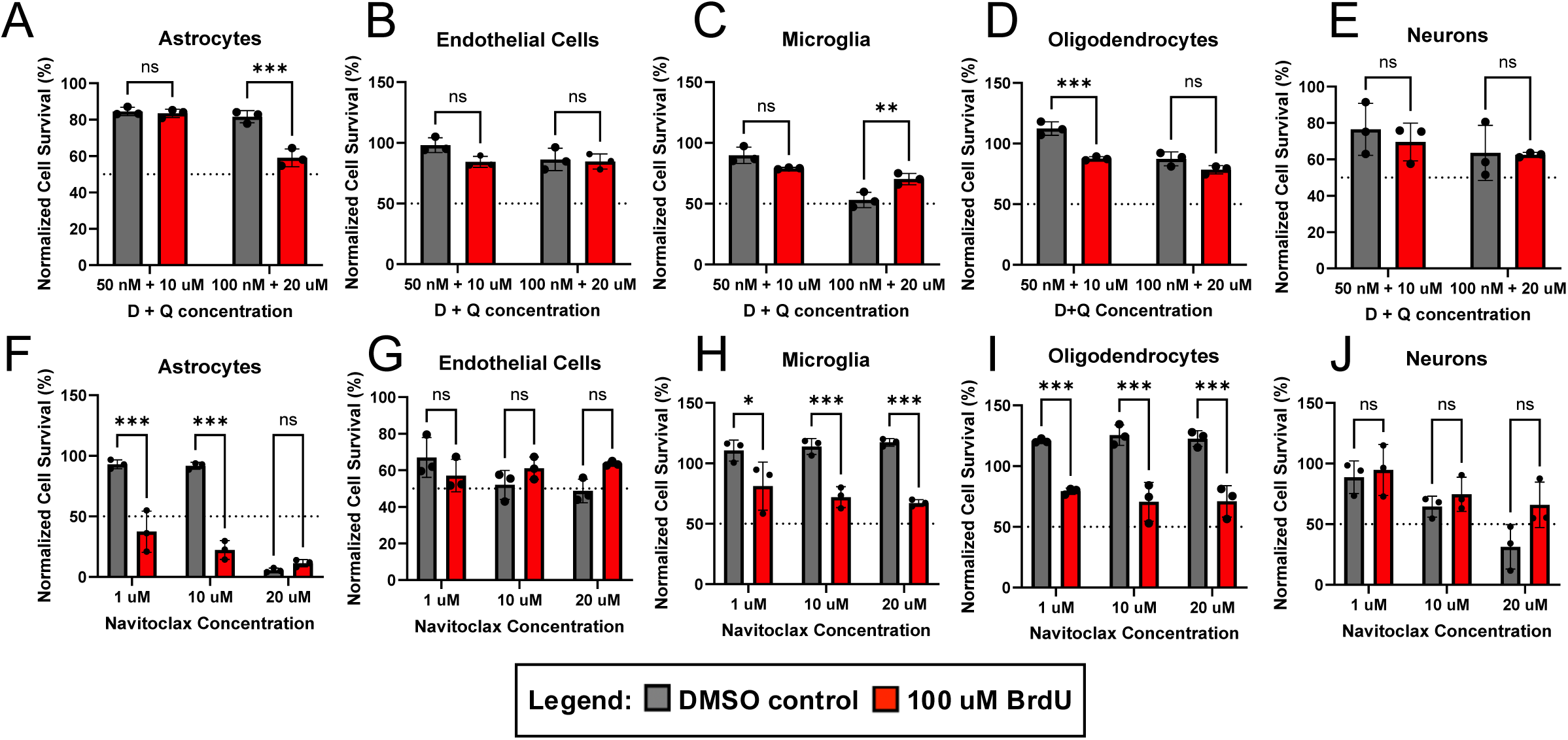
Induced senescent human midbrain cell lines show cell type-specific responses to senolytic treatment. A-E) Normalized percent cell survival of astrocytes (A), endothelial cells (B), microglia (C), oligodendrocytes (D), and neurons (E) following 7-day DMSO (grey) and 100 uM BrdU (red) treatment with 48-hour dasatinib (D) + quercetin (Q) treatment at concentrations of 50 nM D + 10 uM Q and 100 nM D + 20 nM Q (n=3). F-J) Normalized percent cell survival of astrocytes (F), endothelial cells (G), microglia (H), oligodendrocytes (I), and neurons (J) following 7-day DMSO (grey) and 100 uM BrdU (red) treatment with 48-hour navitoclax treatment at concentrations of 1 uM, 10 uM, and 20 uM (n=3). Data analyzed by two-way ANOVA with Šídák’s multiple comparisons test. All graphs show mean with standard deviation (ns, p>0.05, * p<0.05, ** p<0.01, *** p<0.001).

### Principal component analysis reveals a cell type-specific senescence profile

We next set out to determine whether we could isolate a cell type-dependent profile of senescence in an unsupervised manner. We performed a principal component analysis (PCA) as a form of dimensionality reduction on the RNA-seq data from our five human brain cell types (astrocytes, endothelial cells, microglia, oligodendrocytes, and neurons) and treatment groups (DMSO and 100 uM BrdU). The first three principal components in a PCA dataset have been previously described to contain biologically relevant information, whereas higher-order components are typically correlated with irrelevant information or noise (*26, 27*). PCA was able to capture 94% of the explained cumulative variance within the first five principal components, and 75.1% within the first three (Figure 4A). We compared the correlation coefficients across different cell types and treatment groups which showed a strong correlation within cell type groups, suggesting that cells will predominantly maintain their cell identity even following chronic treatment with BrdU (Figure 4B). Based on our phenotypical clustering of both cell type and treatment group, we concluded that PC1 clusters conditions by cell type independent of the treatment group, whereas PC2 continues to maintain cell type identity yet is influenced by treatment (Figure 4C). Strikingly, PC3 is capable of unanimously isolating a senescence signature and separates BrdU samples from DMSO controls, independent of cell types (Figure 4C). When observing the genetic contributions to the PCA loadings from PC3, one can identify that the most enriched positive and negative loadings associated with PC3 are inversely contributing to PC1, or cell type (Figure 4D). Notably, genes such as *SLC7A1, HSP90AB1, PARP4, POMP1*, and *ENO1,* are all observed to be contributing to the senescence phenotype as described by PC3 (Figure 4D). Combining contributions from PC1 and PC3 allows us to isolate a cell type-specific signature of BrdU-induced senescence (Figure 4E). To further characterize these cell type-specific contributions, we directly compared treatment groups through PCA and compared the contribution of those gene loadings across cell types (Figure 4F). Interestingly, the top 10 most highly enriched genes contributing to senescence in each cell type were specific to that cell type yet were never inversely observed in other cell types (with the exception of MYH9 in microglia) (Figure 4G). Altogether, each human brain cell type has its own senescence identity following senescence induction with BrdU treatment. Despite this, one can isolate a senescence signature independent of cell type based on the dimensionality reduction of the transcriptomic data.

**Figure 4:**
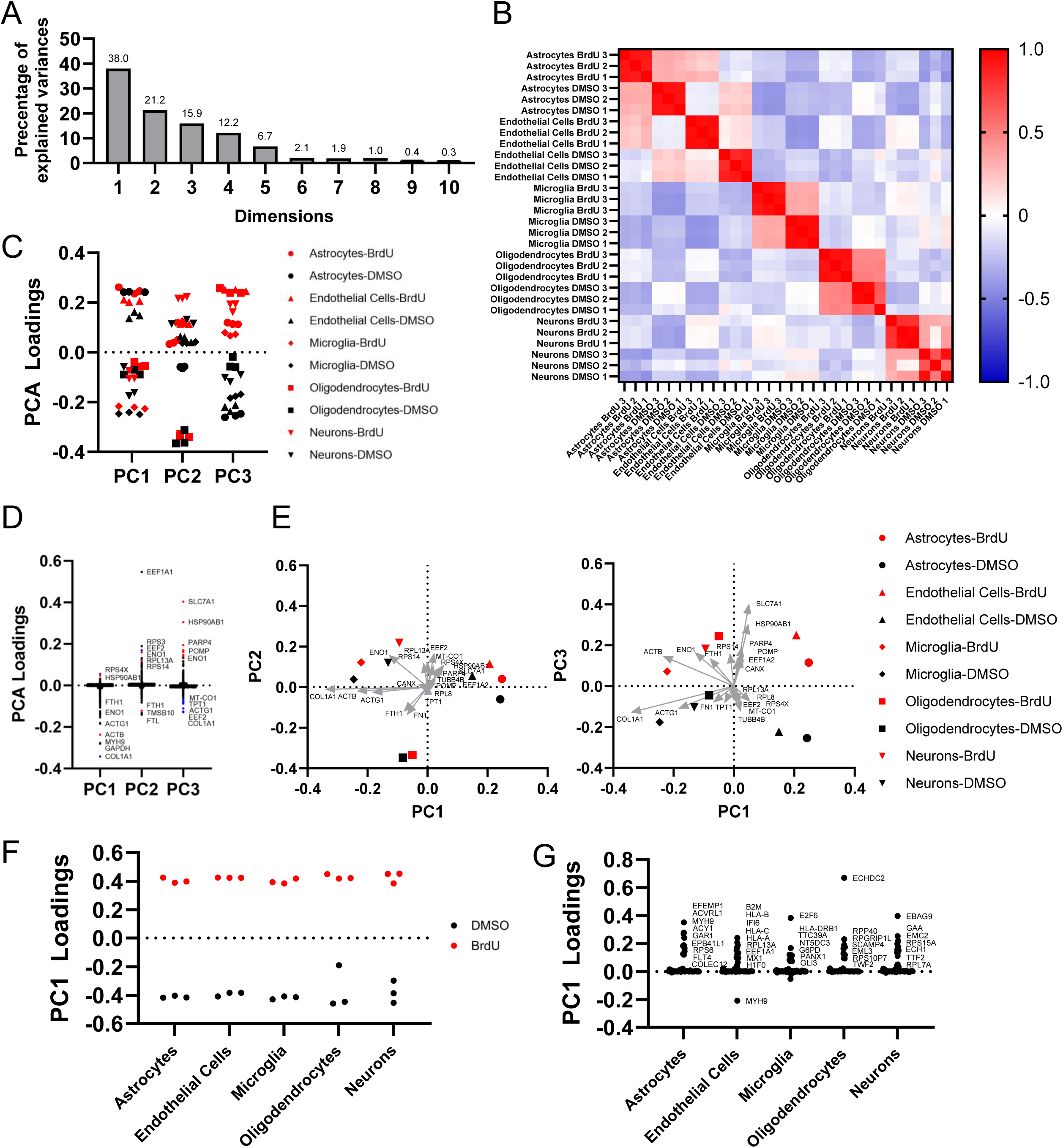
Principal component analysis reveals a cell type-specific senescence profile. A) Percentage of explained variance of each dimension in the principal component analysis of 7-day DMSO control and 100uM BrdU treated human cell lines (astrocytes, endothelial cells, microglia, oligodendrocytes, and neurons). B) Correlation matrix of DMSO control and BrdU treated human cell lines. C) PCA loadings from PC1, PC2, and PC3 of DMSO control (black) and BrdU (red) treated human cell lines. D) PCA loadings of genetic contributions towards PC1, PC2, and PC3 from (C). (Red) and (blue) data points constitute positive and negative loadings from PC3, respectively. E) Two-dimensional graphical visualizations of PC1 and PC2, and PC1 and PC3 from (C) with the top genetic contribution vectors overlayed from PC3. F) PC1 loadings from a PCA directly comparing 7-day treated DMSO control (black) and 100uM BrdU (red) within cell types. G) PCA loadings of the top 10 genetic contributors to PC1 for each cell type plotted across all cell types (50 genes per cell type).

### Regulatory network inference of induced senescent cells allows for the identification of SATRs across cell types

The identification of a common senescence gene expression signature from these senescent human midbrain cell types prompted investigations into a common regulatory gene network. To determine specific transcriptional perturbations, we inferred and reconstructed a transcriptional network utilizing the ARACNe algorithm and a previously established set of human transcription factors (TF) (*17*). Based on this reconstructed transcriptional network, we utilized one- and two-tailed gene set enrichment analysis comparing each treatment group across each cell type to isolate the regulon activity of each TF. We then further normalized the differential enrichment scores (NES) for gene set size and unbiasedly took the top enrichment scores for activated (positive) and repressed (negative) regulators upon senescence induction and found significantly enriched regulators with both common and cell type-specific enrichment (Figure 5A). We observed that transcriptional regulators such as FIZ1, SOX12, THAP5, CHCHD3, and CENPA were the most commonly repressed, whereas MSANTD1, ZNF790, ZBTB40, DPF1, and EGR3 were the most commonly activated (Figure 5A). Other regulators, such as ERF, were significantly repressed but in a cell type-specific manner (Figure 5A). We further compared the differential expression of these regulators across cell types and ranked them based on their average expression (Figure 5B, C). We observed significant differential expression in a subset of regulators, specifically PA2G4, SOX12, CHCHD3, FIZ1, ERF, and TFAP4 (Figure 5B,C). Other regulators, such as MSANTD1 and THAP5, were significantly activated in senescence but were found to not be differentially expressed (Figure 5B,C). From here, we decided to focus on a subset of transcriptional regulators which met two requirements: the transcriptional regulator is both significantly enriched and differentially expressed across cell types. The resulting SATRs were further distinguished by observing the downstream transcriptional targets. We therefore focused on SOX12, CHCHD3, PA2G4, and TFAP4 (Figure 5D). As a comparison to commonly enriched regulators, we also included ERF as a regulator that is significantly enriched in most but not all cell types and is also differentially expressed (Figure 5C,D). We then isolated a list of predicted transcriptional targets and observed that transcriptional regulators such as *CENPA* and *FIZ1*, which are observed to be significantly repressed, are also downstream targets of PA2G4 (Figure 5E). Other notable transcriptional targets are *CDK4* and *STMN1* for PA2G4, *NHEJ1* for SOX12 and CHCHD3, and *CXCL14* for SOX12 (Figure 5E). Enrichr (*28–30*) reaction pathway analysis of these downstream targets highlighted various cell cycle dysregulation pathways for PA2G4 and TFAP4, defects in vitamin metabolism and cholesterol biosynthesis pathways for SOX12, and defects in mitochondrial translation pathways for CHCHD3 (Figure 5F). There were no significant pathways observed for ERF (Figure 5F), which may be a result of the variable transcriptional activity of this regulator across cell types. Overall, we were able to isolate SATRs which were predicted to contribute to the senescence signature following BrdU-induced DNA damage across all five midbrain cell types.

**Figure 5:**
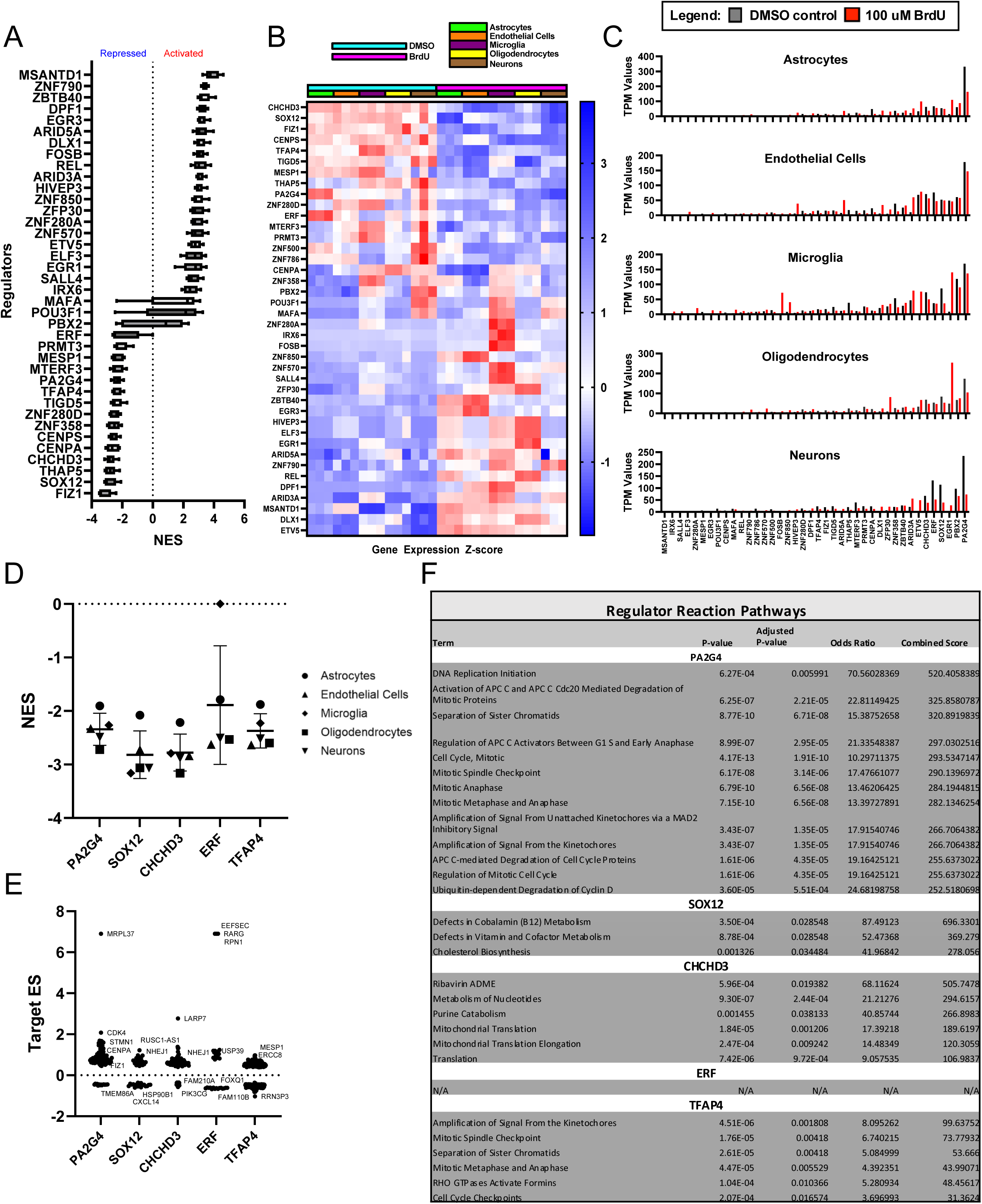
Regulatory network inference of induced senescent cells allows for the identification of SATRs across cell types. A) Average Normalized Enrichment Scores (NES) from the top 10 activated and repressed transcriptional regulators following 100uM BrdU treatment from all 5 cell types. B) Z-score heatmap showing differential gene expression of the 20 transcriptional regulators from (A) of BrdU treatment (pink) as compared to DMSO control (blue) in human cell lines (astrocytes – green, endothelial cells – orange, microglia – purple, oligodendrocytes – yellow, neurons – brown). C) TPM values of the 20 transcriptional regulators from (A) of DMSO control (black) and BrdU (red) treated human cell lines. D) NES values of PA2G4, SOX12, CHCHD3, ERF, and TFAP4 following 100uM BrdU treatment from all 5 cell types. E) Inferred Target Enrichment Scores (ES) from PA2G4, SOX12, CHCHD3, ERF, and TFAP4 transcriptional networks. F) Regulator Reaction Pathways from Enrichr (*28–30*) analysis of inferred PA2G4, SOX12, CHCHD3, ERF, and TFAP4 transcriptional networks (padj<0.05). No significant pathways isolated from ERF transcriptional network.

### Functional characterization of SATRs in human midbrain cell lines

To functionally characterize the selected SATRs, we knocked down (KD) ERF, CHCHD3, SOX12, PA2G4, and TFAP4 in all five midbrain cell types (Figure S6A-E). In addition to their predicted biological pathways (Figure 5F), each of these selected regulators have been previously described in the context of cell cycle dysregulation, apoptosis, and cellular senescence (Figure 6A) (*18, 31–35*). We sought to determine which senescence hallmarks could be recapitulated solely by the repression of the various regulators as compared to our BrdU-induced senescence model. First, we examined the percentage of senescence-associated β-galactosidase (SA β-gal) positive cells following treatment of all five cell types with scrambled control (siSCR) and regulator KD (siRegulator) for each of the selected regulators. KD of ERF, SOX12, and PA2G4 resulted in a significant increase in the percentage of SA β-gal positive astrocytes and KD of ERF led to a significant increase in SA β-gal positive microglia (Figure 6B). We saw no significant changes in SA β-gal positive endothelial cells, oligodendrocytes, or neurons with KD of any of the selected regulators, indicating a cell type-specific regulation by SATRs of the elevated senescence-associated β-gal phenotype. Next, we isolated RNA from siSCR and siRegulator-treated cells for each cell type and ran qPCRs for senescence markers *CDKN1A*, *CDKN2A*, *CDKN2D*, and *LMNB1*. With KD of ERF, we observed a significant senescence-like upregulation of *CDKN1A* only in neurons and downregulation of *LMNB1* in astrocytes and oligodendrocytes (Figure 6C). Next, siRNA KD of CHCHD3 showed significant upregulation of *CDKN1A* in astrocytes and upregulation of *CDKN2A* in neurons with significant downregulation of *LMNB1* only in microglia (Figure 6D). SOX12 KD resulted in the significant downregulation of *LMNB1* in astrocytes and oligodendrocytes (Figure 6E). KD of PA2G4 showed a significant increase in *CDKN1A* in microglia and oligodendrocytes as well as a significant upregulation of *CDKN2A* in microglia, whereas no significant changes in *LMNB1* levels were seen in any cell type (Figure 6F). Lastly, KD of TFAP4 resulted in a significant upregulation of *CDKN1A* in astrocytes, oligodendrocytes, and neurons and an interestingly significant increase of *CDKN2D* levels in microglia (Figure 6G), which mirrored what we saw with BrdU treatment (Figure 1H). Additionally, TFAP4-KD astrocytes, microglia, oligodendrocytes, and neurons all displayed significantly diminished *LMNB1* levels (Figure 6G). Based on the functional characterization of our selected SATRs, we can conclude that each regulator contributes to a cell type-specific signature of subsets of cellular senescence phenotypes. Importantly, no single regulator was able to fully recapitulate all the tested hallmarks in all cell types, suggesting a pathway-specific regulation of senescence hallmarks.

**Figure 6:**
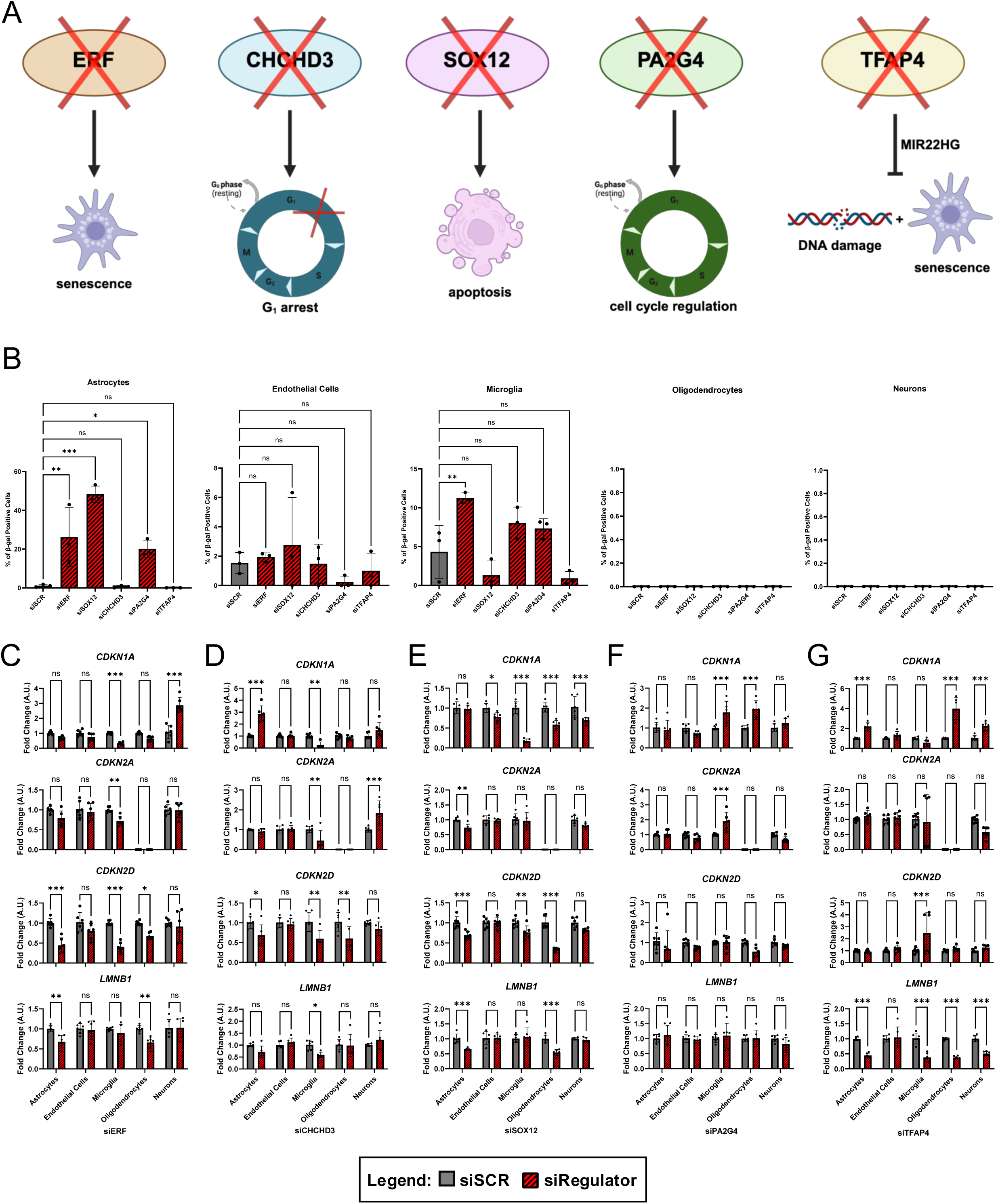
Functional characterization of SATRs in human midbrain cell lines. A) Schematic depicting selected transcriptional regulators (left-right) ERF, CHCHD3, SOX12, PA2G4, and TFAP4 and known associations of their knockdown (KD) in the context of cell cycle dysregulation, apoptosis, and senescence (created with BioRender). B) Quantification of percentage of senescence-associated β-galactosidase (β-gal) positive cells in astrocytes, endothelial cells, microglia, oligodendrocytes, and neurons following transcriptional regulator KD (red) compared to control (grey) (n=3). C-G) Quantitative PCR of siSCR (grey) and siRegulator (red) (ERF (C), CHCHD3 (D), SOX12 (E), PA2G4 (F), and TFAP4 (G)) treated astrocytes, endothelial cells, microglia, oligodendrocytes, and neurons of senescence markers *CDKN1A*, *CDKN2A*, *CDKN2D*, and *LMNB1* (top-bottom, n=5-6). Data analyzed by two-way ANOVA with Šídák’s multiple comparisons test. All graphs show mean with standard deviation (ns, p>0.05, * p<0.05, ** p<0.01, *** p<0.001).

### Knockdown of SATR TFAP4 results in senescence phenotypes in human midbrain cell lines

TFAP4 has been well-established as a regulator of p21/p16 expression via *MIR22HG* (*36*), which is a miRNA that we have shown can regulate dopaminergic neuron senescence via PD risk factor *GBA* (*37*). First, PCA of TFAP4 transcriptional targets identified through gene set enrichment analysis reveals a senescence-associated signature in all cell types (Figure 7A, B). Based both on our results showing that KD of TFAP4 led to gene expression changes characteristic of a senescence phenotype and previous findings that loss of TFAP4 and associated increases in *MIR22HG* resulted in senescence phenotypes (*18*), we chose to focus on TFAP4 to further characterize senescence hallmarks associated with siRNA KD (Figure 7C, D). First, we looked at *MIR22HG* levels following TFAP4 KD in all five midbrain cell types and observed a significant increase in microglia, oligodendrocytes, and neurons with TFAP4 KD (Figure 7E). We immunostained siSCR and siTFAP4 treated cells for lamin B1 and saw a significant decrease in lamin B1 intensity with TFAP4 KD in microglia and oligodendrocytes (Figure 7F, S7A-B). Interestingly, we quantified nuclear area and only saw a significant increase in TFAP4 KD astrocytes (Figure 7G). Based on our observations of mitochondrial and lysosomal dysfunction across cell types following BrdU treatment (Figure 2), we used Mitotracker Red CMXRos and Lysotracker Deep Red staining in TFAP4 KD cells to investigate whether a loss of TFAP4 led to a disruption or lack of turnover of mitochondria and/or lysosomes in any cell type. We observed a significant increase in the number of mitochondria per cell (Figure 7H, S7A) and in the number of lysosomes per cell (Figure 7I, S7B) in TFAP4 KD astrocytes as compared to control. We further characterized siTFAP4 cells for γH2AX foci as a molecular marker of DNA damage and showed that astrocytes displayed an increased number of γH2AX foci/cell following TFAP4 KD (Figure 7J, S7A). Taken together, these results further demonstrate that targeting a SATR such as TFAP4 can recapitulate senescence hallmarks in a cell type-specific manner.

**Figure 7:**
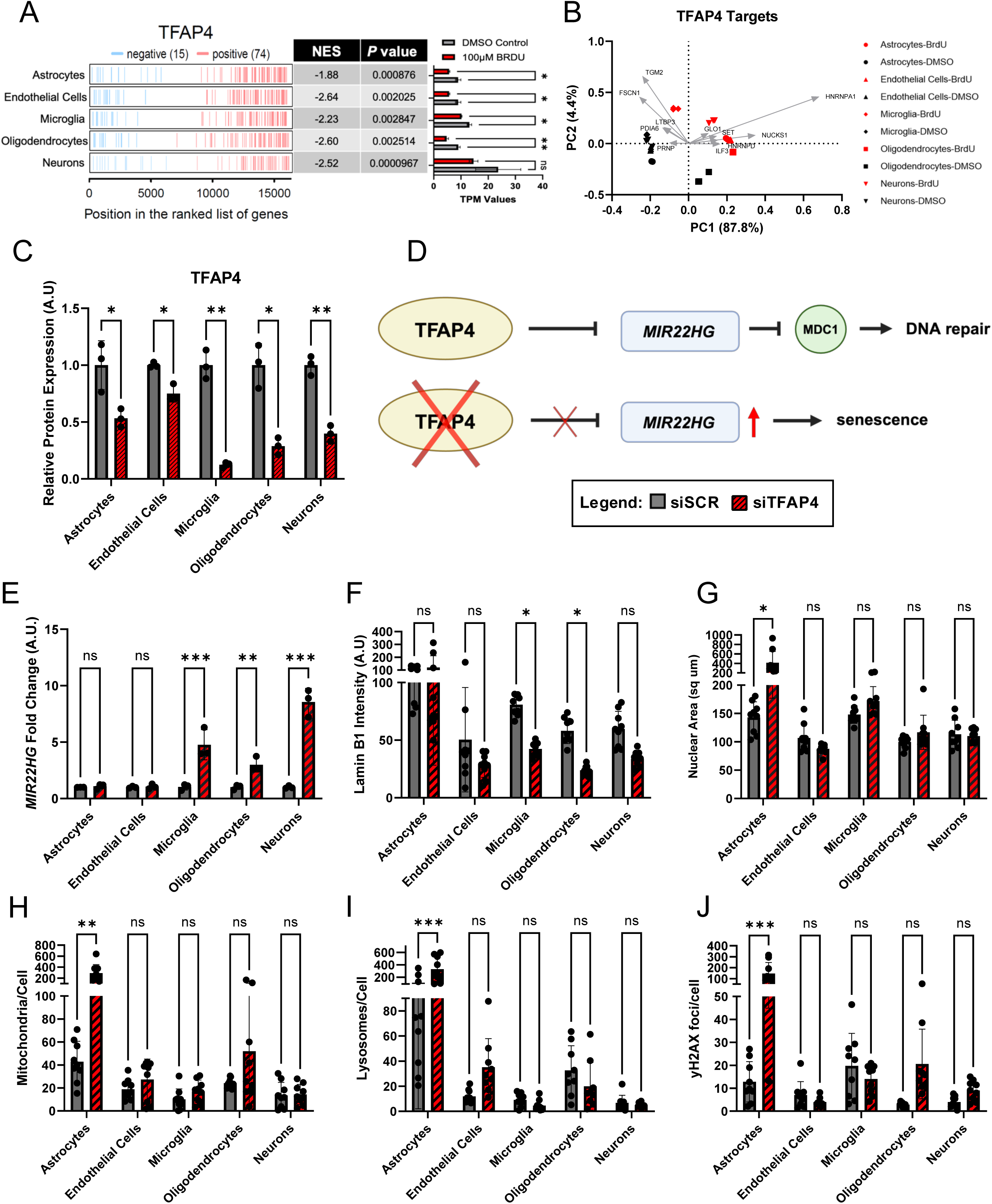
Knockdown of SATR TFAP4 results in senescence phenotypes in human midbrain cell lines. A) Two-tailed Gene Set Enrichment Analysis of TFAP4 from an inferred transcriptional network from 100uM BrdU treated human cell lines (astrocytes, endothelial cells, microglia, oligodendrocytes, and neurons). Left - Activated (red) or inhibit ed (blue) target elements and their position in the ranked list of genes as identified by log2FoldChange. Center - Normalized Enrichment Scores (NES) identified by two-tail gene set enrichment analysis and associated p-value. Right - Transcripts per Million (TPM) values of indicated regulons in DMSO and BrdU treated cells. B) Two-dimensional graphical visualizations based on PCA of TFAP4 transcriptional targets representing PC1 and PC2 with the top genetic contribution vectors overlayed from PC1. C) Normalized TFAP4 protein expression relative to β-Actin from Western blots shown in Figure S6 in siSCR (grey) and TFAP4 KD (red) astrocytes, endothelial cells, microglia, oligodendrocytes, and neurons (n=3). D) Schematic depicting known regulation of *MIR22HG* by TFAP4 based on previously published data (*18*) where loss of TFAP4 leads to increased *MIR22HG* levels and subsequently results in a senescence phenotype (created with BioRender). E) Quantitative PCR of siSCR (grey) and siTFAP4 (red) treated astrocytes, endothelial cells, microglia, oligodendrocytes, and neurons of normalized *MIR22HG* levels (n=3). F) Quantification of immunofluorescence staining of lamin B1 in siSCR (grey) and siTFAP4 (red) treated human cell lines (n=9). G) Quantification of nuclear area based on immunofluorescence staining of lamin B1 (nuclear size (sq um)) in siSCR (grey) and siTFAP4 (red) treated cells (n=9). H) Quantification of Mitotracker Red CMX Ros staining in siSCR (grey) and siTFAP4 (red) treated cells (n=9). I) Quantification of Lysotracker Deep Red staining in TFAP4 KD (red) as compared to controls (grey) (n=9). J) Quantification of immunofluorescence staining of DNA damage marker γH2AX in siSCR (grey) and siTFAP4 (red) treated human cell lines (n=9). Data analyzed by two-way ANOVA with Šídák’s multiple comparisons test. All graphs show mean with standard deviation (ns, p>0.05, * p<0.05, ** p<0.01, *** p<0.001).

## Discussion

Through the investigations presented here, we have provided a detailed characterization of an *in vitro* senescence profile in various midbrain cell types which further elucidated a cell type-specific profile of senescence. Chronic treatment with BrdU in astrocyte, endothelial, microglial, oligodendrocyte, and neuronal cell lines was able to induce cellular senescence in all five cell types based on the expression of several established hallmarks of senescence. However, we highlight that although the same senescence-inducing stressor was used, each cell type expressed its own senescence profile based on variability in the hallmarks observed as well as the degree of their expression. Senescence has been classically defined by key phenotypes, including proliferative arrest, increased SA β-gal activity, elevated levels of p21/p16/p19, SASP expression, mitochondrial and/or lysosomal alterations, presence of SAHF, and loss of lamin B1 (*4–7*). Defining a senescent cell *in vivo* based on observed phenotypes has been recently summarized to specifically include (1) quantification of p21 and/or p16 mRNA or protein levels, (2) verification with additional markers such as *LMNB1* downregulation, SASP factors, SA β-gal, and DNA damage, (3) targeted removal or elimination of the cells with senomorphics or senolytics, and (4) confirmation that reduction of the senescent cells is sufficient to ameliorate a given phenotype (*8*).

In the data presented here, we observed classical senescence hallmarks including elevated SA β-gal (Figure 1B) and proliferation arrest (Figure 1C) in all five cell types following BrdU treatment. However, we observed differences across cell types that highlighted various interesting patterns in the expression of classically described senescence hallmarks. All cell types other than microglia displayed both BrdU-dependent increases in the nuclear area (Figure 1E) and in *CDKN1A* (p21) mRNA levels (Figure 1H). Interestingly, senescent microglia displayed elevated *CDKN2D* (p19) levels, which is in line with reports of p19-positive microglia from P301S tau mice in the context of AD (*38*). Interestingly, all other cell types also showed a significant upregulation of p19 (*CDKN2D)* expression following BrdU treatment which emphasizes its potential as a more representative CDK inhibitor driving senescence in this context. Additionally, classical hallmark p21 protein expression (Figure 1F, S1D) was only significantly increased in astrocytes, endothelial cells, and oligodendrocytes following BrdU treatment, which aligned with changes in cathepsin D activity (Figure 2F) and accumulation of mitochondria (Figure 2D). Taken together, our data emphasizes the importance of assaying multiple phenotypes when defining a senescent cell, considering not only the cell type under investigation but also the context in which senescence is being induced and the stressor(s) being manipulated. Importantly, a better understanding of these cell type-specific profiles of senescence has the potential to inform future *in vivo* senescence investigations, whereby detection of senescence phenotypes in each cell type may be made easier by a high throughput characterization of different cell types across various contexts. Furthermore, the identification of cell type-specific pathways could lead to the development of novel senolytics and/or senomorphics.

In the context of BrdU treatment, it is important to note that comparisons of senescence hallmarks between cell types may differ based on varied baseline levels and the timepoint at which we are measuring the expression of these hallmarks (i.e. early- vs. late-stage senescence). One example of this can be seen with our observed *CDKN2A* (p16) expression levels, where p16 was not detected in oligodendrocytes and was actually decreased with BrdU treatment among other cell types (Figure 1H). Additionally, our investigations into *CDKN1A* (p21) showed variability based on the method of detection (immunostaining (Figure 1F), RNAseq (Figure 1H), and Western blotting (Figure S2D)), but also showed cell type-dependent differences. Both control and BrdU-treated neurons expressed very low levels of p21 and inverse levels of p21 were observed in BrdU-treated microglia. As we demonstrate in the context of BrdU treatment in the midbrain cell lines used, neither p16 nor p21 is a universal marker of senescence across cell types, at least 7-days following BrdU-induced DNA damage. Thus, p21/p16 alone is not sufficient to define the senescence phenotype across the five cell types, further highlighting the importance of assaying various senescence hallmarks. In comparison, markers such as senescence-associated β-gal, proliferation arrest, and increased nuclear size and/or decreased *LMNB1* were more universally observed across cell types. This allowed for not only more direct comparisons across cell types but also defines them as more robust and comprehensive markers of a senescence phenotype following BrdU treatment.

In the context of mitochondrial and lysosomal dysfunction, we similarly observed variability amongst the senescence hallmarks we assayed. When determining the oxygen consumption rate (OCR), all cell types showed a general downward trend following senescence induction, yet when OCR was compared across cell types, we observed large variations. Specifically, astrocytes showed a range in basal respiration of 5-60 pmol/min, whereas oligodendrocytes showed a range of 50-446 pmol/min. We also investigated cell type-specific responsiveness to well-established senolytic compounds D+Q and navitoclax (*24, 25*) and observed high sensitivity of senescent oligodendrocytes and astrocytes to both D+Q and navitoclax (Figure 3), with BrdU-treated microglia also showing navitoclax sensitivity. The differences we observed in the ability of the two well-characterized senolytic cocktails to eliminate BrdU-induced senescent cell lines offers an interesting perspective in the future development of senolytic drugs, which should further consider varying responsiveness based on the senescence profile of different cell types.

Following our central theme of cell type-dependent expression of senescence phenotypes, the investigations into unbiasedly isolating a senescence profile of midbrain cell types through PCA emphasizes our ability to isolate both unique and common senescence signatures from different cell types (Figure 4). At first glance, comparing all cell types through PC1 reveals clusters based on cell identity, highlighting that induction of senescence does not alter the primary transcriptional signature of a given cell type. This is further alluded to by directly comparing the genes that contribute to the senescence-phenotype following BrdU treatment within a cell type, which does not positively contribute to this signature in other cell types (Figure 4F,G). However, further comparisons across cell types through PC3 reveal that there is also a common senescence signature (Figure 4C-E). Therefore, a common and cell type-specific senescence profile predicts upstream master regulators of transcription (SATRs), which can functionally regulate observed senescence responses both within and across these different midbrain cell types (*39*). Our investigations into SATRs such as ERF, CHCHD3, SOX12, PA2G4, and TFAP4 highlight that each regulator can recapitulate some hallmarks of senescence, but not all, and does so in a cell type-dependent manner. For example, the knockdown of PA2G4 (EBP1), which has been previously implicated in the cell cycle and AD (*40, 41*), significantly induces β-gal expression only in astrocytes, resulting in the upregulation of *CDKN1A* in both microglia and oligodendrocytes and *CDKN2A* in microglia, but fails to induce any changes in *CDKN2D* and *LMNB1*. Therefore, examining only β-gal expression or p21/p16 upregulation as a senescence-associated response in a given cell type could highlight PA2G4 as a regulator in that context, but only when comparing all of the phenotypes assayed here can we see that PA2G4 only induces specific responses in a cell type-dependent manner.

We chose to focus on the SATR TFAP4 based partially on its previous association with senescence in the context of directly downregulating p16 and p21 in fibroblasts (*36*) and induction based on varied c-Myc levels (*42*). Additionally, and most interestingly, TFAP4 has been shown to suppress DNA damage and senescence via repression of *MIR22HG* (miR-22-3p) (*18*), which we have recently shown to be connected to PD risk factors SATB1 and GBA in the context of lipid-induced DA neuron senescence (*37, 43*). Specifically, our functional characterization of senescence hallmarks in TFAP4 KD cells revealed an upregulation of *MIR22HG* only in microglia, oligodendrocytes, and neurons (Figure 7D), which is in line with previous reports of TFAP4- dependent *MIR22HG* regulation in cancer cells (*18*). Interestingly, we also observed a significant decrease in lamin B1 protein levels in microglia and oligodendrocytes and a trending decrease in neurons (p=0.11, Figure 7E), indicating that the TFAP4-*MIR22HG* pathway may specifically mediate the lamin B1 phenotype in senescent cells. Our experiments using BrdU treatment did not lead to the same pattern of hallmark expression in these three cell types in any of the phenotypes we assessed (Figures 1-3) which suggests that a cell type-dependent senescence phenotype can also be dependent on the method of senescence induction and may be regulated by the concerted action of a subset of SATRs. When we further characterized additional senescence hallmarks in TFAP4 KD cells, such as DNA damage, mitochondrial and lysosomal dysfunction, and nuclear size, we only observed significant senescence-associated changes in astrocytes following TFAP4 KD (Figure 7F-I). In addition to our investigations into specific senescence phenotypes that may be modulated by these upstream SATRs, it is also possible to manipulate these inferred regulators in the context of senolytic drug development to specifically target certain senescent cell types *in vitro* and *in vivo*.

Lastly, our characterization of a cell type-specific senescence profile in these midbrain cell types informs senescence in the context of neurodegeneration and specifically PD, where DA neurons of the SNpc have been shown to become senescent (*11*). Importantly, the cell types investigated have all been shown to be capable of becoming senescent in the context of aging, neurodegeneration, or even AD and PD specifically (*12, 38, 44–46*). Taken together, this leads to the interesting idea that these five different midbrain cell types may have the potential to become senescent via the spreading of senescence (via SASP). Thus, it would be important to have defined a cell type-specific profile of senescence to better understand the mechanisms by which senescence may spread between distinct cell types both *in vitro* and *in vivo* where a given cell type could be successfully defined as senescent. Based on our results shown here, we have defined and thoroughly characterized this cell type-specific senescence phenotype as well as various interesting functional regulators of senescence, here called SATRs, that were inferred based on our unbiased, systematic approach of inducing a senescence phenotype in various midbrain cell lines with BrdU-induced DNA damage. Finally, the SATRs identified here could potentially be used as cell type-specific drug targets for either developing novel senolytics, senescence biomarkers, or senescence inducers for cancer treatment.

## Methods

### HMC3 (microglia) cell culture

HMC3 microglial cells (ATCC) were cultured according to the vendor’s protocol. In brief, cells were grown at 37°C in 5% CO_2_. The used medium is ATCC-formulated Eagle’s Minimum Essential Medium (Catalog No. 30-2003) with 1% Pen/Strep and fetal bovine serum to a final concentration of 10% added. Cells were passaged every 2-3 days with Trypsin-EDTA (0.25%, ThermoFisher Catalog No. 25200072).

### SK-N-MC (neurons) cell culture

SK-N-MC cells (ATCC) were cultured according to the vendor’s protocol. In brief, cells were grown at 37°C in 5% CO_2_. The used medium is ATCC-formulated Eagle’s Minimum Essential Medium (Catalog No. 30-2003) with 1% Pen/Strep and fetal bovine serum to a final concentration of 10% added. Cells were passaged every 2-3 days with Trypsin-EDTA (0.25%, ThermoFisher Catalog No. 25200072).

### HBEC-5i (endothelial cells) cell culture

HBEC-5i microvascular endothelial cells (ATCC) were cultured according to the vendor’s protocol. In brief, cells were grown at 37°C in 5% CO_2_. The used medium is ATCC DMEM:F12 (Catalog No. 30-2006) with a Microvascular Growth Supplement (ThermoFisher, Catalog No. S00525) added, as well as 1% Pen/Strep and fetal bovine serum to a final concentration of 10%. Cells were passaged every 2-3 days with Trypsin-EDTA (0.25%, ThermoFisher Catalog No. 25200072) and were plated on vessels coated with 0.1% gelatin (Sigma, Catalog No. G1393) in H_2_O for 45 minutes at 37°C.

### HOG (oligodendrocytes) cell culture

HOG cells (Sigma) were cultured according to the vendor’s protocol. In brief, cells were grown at 37°C in 5% CO_2_. The used medium is ATCC DMEM (Catalog No. 30-2002) with 1% Pen/Strep and fetal bovine serum to a final concentration of 10% added. Cells were passaged every 2-3 days with StemPro Accutase (Thermo Fisher, Catalog No. A1110501).

### SVG-A (astrocytes) cell culture

The SVG-A cell line was kindly provided by Dr. Walter Atwood (Brown University, Providence, RI, USA). These cells were cultured as previously reported at 37°C in 5% CO_2_ in ATCC DMEM (Catalog No. 30-2002) with 1% Pen/Strep and fetal bovine serum to a final concentration of 10% (*47*). Cells were passaged every 2-3 days with Trypsin-EDTA (0.25%, Thermo Fisher, Catalog No. 25200072).

### BrdU treatment of human cell lines

Treatment for 7 days with 100 uM 5-bromo-2’-deoxyuridine (BrdU) (Sigma, Catalog No. B5002) dissolved in DMSO (Sigma, Catalog No. D2650) was used to induce senescence in the human cell lines. A 0.1% DMSO solution in the appropriate cell line growth media was used as a control. Previous reports have shown that BrdU treatment activates a DNA damage response and can induce a senescence phenotype in mammalian cells, regardless of cell type (*15, 16, 19, 48–50*).

### Senescence-associated β-galactosidase (SA-β-gal) assay

To test whether BrdU-treated cell lines displayed a lysosomal dysfunction phenotype consistent with entering a state of cellular senescence, a senescence-associated β-galactosidase (SA-β-gal) assay (Cell Signaling Technology, Catalog No. 9860) was performed according to the manufacturer’s protocol. In brief, after the respective 7-day treatment, cells were fixed and incubated over night at 37°C with X-gal as substrate for the SA-β-galactosidase. The following day, the substrate solution was removed, 70% glycerol (in PBS) was added to the fixed cells, and wells were imaged with a microscope (Accu-Scope, EXI-410, Skye Color Camera). Analysis and quantification of images was performed using Fiji and statistical analyses were performed using Prism (GraphPad).

### RNA isolation and RNAseq of DMSO- and BrdU-treated human cell lines

RNA was isolated from cell lines following 7-day treatment with DMSO or 100 uM BrdU using the RNeasy Plus Mini Kit (QIAGEN, Catalog No. 74134) and QIAshredder (QIAGEN, Catalog No. 79656) and 500 ng of RNA for each sample was sent to Azenta for bulk RNAseq. In brief, sample quality control and determination of concentration was performed using TapeStation Analysis by Azenta, followed by library preparation and sequencing. Computational analysis included in their standard data analysis package was used for data interpretation. Target genes were pulled from known lists of genes associated with senescence, senescence-associated secretory phenotype (SASP) (*51*), mitochondrial and lysosomal function (*21, 22*). Samples used in heatmaps were normalized by z-score.

### Live imaging and growth curve analysis

Following 7-day treatment with DMSO or 100 uM BrdU, cells were passaged and seeded on 24-well plates (Sarstedt, Catalog No. NC0984607) at equal densities amongst conditions. Using the Agilent LionheartFX Live Imaging System, three regions of interest per well were imaged at 20x every 15 minutes for 48 hours. Cell growth was standardized by starting cell number.

### Immunofluorescent staining of senescence markers

12 mm round glass coverslips (VWR, Catalog No. 73605-514) were washed in 70% ethanol, rinsed with PBS, and allowed to air dry for 60 minutes under UV light to sterilize. Coverslips were then placed into each well of a 24-well plate, 500 uL of 0.1% gelatin in H_2_O was added to each well, and the plate was incubated at 37°C for at least 30 minutes. The gelatin solution was then removed, and cells were plated onto coverslips for 7-day treatment with DMSO or 100 uM BrdU. Following treatment, cells were chemically fixed with 4% PFA in PBS for 20 minutes at room temperature. Cells were then permeabilized with 0.2% TritonX-100 in PBS for 10 minutes at room temperature followed by 3 5-minute PBS washes and then blocking in 5% NDS/PBS/0.1% Tween20 at room temperature for 30 minutes. The cells were then incubated with the respective primary antibody in 5% NDS/PBS/0.1% Tween20 overnight at 4°C. Primary antibodies used include γH2AX (pS139) (Abcam, Catalog No. ab303656), lamin B1 (Abcam, Catalog No. ab16048), and p21 (Abcam, Catalog No. ab109520). The following day, cells were washed 4 times for 5 minutes each in 0.1% Tween20-PBS and then incubated with the respective secondary antibodies in 5% NDS/PBS/0.1% Tween20 for 2 hours in the dark at room temperature. Secondary antibodies used include Alexa Fluor 488 d@ms IgG (Invitrogen, Catalog No. A21202), Alexa Fluor 488 d@Rb (Invitrogen, Catalog No. A21206), Alexa Fluor 546 d@Rb IgG (Invitrogen, Catalog No. A10040), and Alexa Fluor 647 d@Rb (Invitrogen, Catalog No. A31573). The cells were then rinsed again 4 times for 5 minutes each in 0.1% Tween20-PBS and then coverslips with cells were mounted onto slides (top down) with 20 uL of DAPI Fluoromount-G (Southern Biotech, Catalog No. 0100-20). Slides were then cured in the dark at room temperature overnight. Confocal images were taken with an Olympus FV3000 Laser Scanning Confocal Microscope. Laser settings (laser strength, gain, and offset) and magnification were maintained across treatment groups. Airyscan imaging was performed using Zeiss LSM 800. Images were taken at 3X zoom with 0.2 AU pinhole size and high resolution Airyscan post-processing. All other settings were optimized by Airyscan digital processing recommendations.

### Fluorescent dyes for imaging mitochondria and lysosomes

For imaging of mitochondria and lysosomes, cells were plated on sterile 12 mm round glass coverslips (as previously described) and exposed to 7-day treatment with either DMSO or 100 uM BrdU. Following treatment, cells were washed 1x with Dulbecco’s PBS without calcium and magnesium (ThermoFisher, Catalog No. 14190094) and then incubated for 15 minutes at 37°C in either 500 nM Mitotracker Red CMXRos (Invitrogen, Catalog No. M7512) in PBS or 1 uM Lysotracker Deep Red (Invitrogen, Catalog No. L12492) in PBS. Following treatment, cells were chemically fixed, permeabilized, washed, blocked, and stained as described above for additional senescence markers. Confocal images were taken with an Olympus FV3000 Laser Scanning Confocal Microscope. Laser settings (laser strength, gain, and offset) and magnification were maintained across treatment groups. Post-processing of images was performed by ImageJ and Cell Profiler as described below.

### Image analyses

Images were analyzed in bulk through Cell profiler. Z-projections were taken from each image by maximum intensity and then separated by fluorophore. Nuclei were identified using DAPI staining. Mitochondria and lysosomal counts were analyzed by applying a size and intensity threshold in Cell Profiler utilizing control samples. Counts per image were then standardized by the number of cells to give an average number of objects per cell. γH2AX foci were analyzed by applying a size and intensity threshold in Cell Profiler utilizing control samples. These foci were then associated with each nucleus. Foci per nuclei were then averaged by the number of cells per image. p21 and lamin B1 intensity were extracted from DAPI labeled nuclei.

### Western blotting for p21

Western blotting for p21 was performed using the Licor-Odyssey Scanner. In short, protein lysates were derived from cell cultures using RIPA buffer (Thermo Scientific, Catalog No. 89900) containing both protease (Roche, Catalog No. 11836170001) and phosphatase (Roche, Catalog No. 4906845001) inhibitors. To determine the protein concentration of each sample, a Pierce BCA assay (Thermo Scientific, Catalog No. PI23225) was performed and quantified using a SpectraMax iD3 (Molecular Devices) plate reader. Equal quantities (10 μg) of protein were boiled in 4X Tris-Glycine-SDS sample buffer (Biorad, Catalog No. 1610747) at 95°C for 5 min and separated using 4%–15% Tris-glycine (Biorad, Catalog No. 4568084). Proteins were transferred to PVDF membranes (Biorad, Catalog No. 1620177) using a semi-dry blotting method. Blots were then blocked in Intercept TBS Blocking Buffer (Licor, Catalog No. 927-60001) for 1 hour at room temperature and subsequently incubated with the respective primary antibody overnight at 4°C in TBS Blocking Buffer supplemented with 0.1% Tween-20 (Biorad, Catalog No. 1610781). The following primary antibodies were used: p21 Rabbit Monoclonal Antibody (Thermo Scientific, Catalog No. MA5-14949, 1:1,000), β-Actin mouse monoclonal IgG1 (Sigma, Catalog No. A2228). Primary antibody was detected with 1 hour incubation at room temperature in TBS Blocking Buffer supplemented with 0.1% Tween-20. The following secondary antibodies were used: IRDye 800CW Goat anti-Rabbit IgG (Licor, Catalog No. 926-32211), IRDye 680RD Goat anti-Mouse IgG (Licor, Catalog No. 926-68070). An Odyssey Classic Scanner (Licor) was used to visualize protein bands, which were subsequently quantified with ImageJ software and normalized to the corresponding β-Actin bands.

### Seahorse XF Mito Stress Test

To measure cellular respiration as a readout for cellular mitochondrial activity, we applied the Seahorse FX Mito Stress Test and Seahorse XFe96 Analyzer (Agilent) according to the manufacturer’s protocol. In brief, cells were plated on a 96-well plate (Sarstedt, Catalog No. NC0506179) and were treated for 7 days with DMSO or 100 uM BrdU as previously described. The following day, cells were washed with XF Cell Mito Stress Test Assay medium (Agilent) and incubated for one hour in a CO_2_-free incubator at 37°C. During the experimental measurement, cells were challenged with 1.0 µM Oligomycin (Port A), 0.5 µM FCCP (Port B) and 0.5 µM Rotenone/Antimycin A (Port C). After the Seahorse run, results were normalized on the protein concentration of every individual well based on a BCA Assay (ThermoFisher, Catalog No. 23225) and a SpectraMax iD3 (Molecular Devices) plate reader. Following normalization, Seahorse results were analyzed using the Wave software (Agilent).

### Cathepsin D enzymatic activity assay

Cathepsin D enzymatic activity was assessed using the Cathepsin D Activity Fluorometric Assay Kit (BioVision, Catalog No. K143-100) according to the manufacturer’s protocol. In brief, cells were treated for 7 days with DMSO or 100 uM BrdU as previously described and were then counted using an EVE Automated Cell Counter (NanoEntek) to normalize cell number for use with the Cathepsin D assay kit. Lysed cells were incubated with master assay mix for 2 hours at 37°C and samples were read with a SpectraMax iD3 (Molecular Devices) plate reader at 328 nm excitation and 460 nm emission.

### Principal component analysis

Principal component analysis (PCA) was performed using the FactoMineR package in R(*52*). In short, RNA-seq expression data was normalized by z-score for each gene list. A correlation matrix was formed for each sample condition using the built in cor() function from the normalized data set. The function princomp() was used to isolate the principal component analysis loadings from the correlation matrix. fviz_eig, fviz_pca_var, and fviz_cos2 were used to visualize the data. The graphs were constructed in GraphPad Prism.

### Transcriptional regulator analysis

The transcriptional regulatory networks were inferred using the Bioconductor RTN package in R (*17, 53*). A list of known human transcription factors was obtained (*54*). In short, the reconstruction of a transcriptional regulatory network (TRN) was inferred from RNA-seq data collected from all 5 cell types (astrocytes, endothelial cells, microglia, oligodendrocytes, and neurons) and from both treatment conditions (7-day DMSO or 100 uM BrdU). Interactions between regulators (TFs) and potential targets were computed based on mutual information (MI), removing unstable interactions by bootstrapping, and then the ARACNe algorithm was applied to remove the weakest interaction between two TFs and a common target gene. From this TRN, we used a two-tailed gene set enrichment analysis with 1000 permutations to assess the regulon-target gene interaction, separating them into a positive or negative targets using Pearson’s correlation and assessing the distribution of the targets across a log2FoldChange ranked gene list. An Enrichment Score (ES) was calculated by walking down the ranked list of target genes and increasing the running sum statistic when encountering a gene in the set and decreasing the statistic when encountering a gene not in the set. The diffferential Enrichment score (dES=posES-negES) was normalized to the mean enrichment score of random samples of the same gene set size (NES). P-values were calculated using log-rank statistics. The top 10 positively and negatively enriched regulons were isolated for each cell type and then further sorted by baseline expression.

### siRNA knockdown of selected transcriptional regulators

Transfection of Silencer Select Pre-Designed Human siRNAs for SOX12 (Life Technologies, Catalog No. 4392420 (s13315)), ERF (Life Technologies, Catalog No. 4390824 (s4808)), TFAP4 (Life Technologies, Catalog No. 4392420 (s14014)), PA2G4 (Life Technologies, Catalog No. 4390824 (s9978)), and CHCHD3 (Life Technologies, Catalog No. 4392420 (s29771)) was performed using Lipofectamine RNAiMAX Transfection Reagent (Thermo Fisher, Catalog No. 13778150) according to the manufacturer’s protocol. In brief, cells were incubated with 5 pmol siRNA per well of a 24-well plate for 48-hours, followed by a fresh media change. Cells were re-transfected with an additional 5 pmol siRNA per well of a 24-well plate 72-hours after the initial transfection and again incubated for 48-hours, with a final media change 120-hours after the initial transfection. Cells were harvested for experiments 7-days after the initial siRNA transfection. Control transfections were performed using a scrambled control (SCR).

### Western blotting for transcriptional regulators

Western blotting was performed as previously described (*37*). In short, protein lysates were derived from cell cultures using RIPA buffer (Thermo Scientific, Catalog No. 89900) containing both protease (Roche, Catalog No. 11836170001) and phosphatase (Roche, Catalog No. 4906845001) inhibitors. To determine the protein concentration of each sample, a Pierce BCA assay (Thermo Scientific, Catalog No. PI23225) was performed and quantified using a SpectraMax iD3 (Molecular Devices) plate reader. Equal quantities (10 μg) of protein were boiled in 2X Tris-Glycine-SDS sample buffer (Thermo Scientific, Catalog No. LC2676) at 95°C for 5 min and separated using 4%–20% Tris-glycine (Invitrogen, Catalog No. XP04200). Proteins were transferred to nitrocellulose membranes (Thermo Scientific, Catalog No. 88018) using a wet blotting method. Blots were then blocked with 5% BSA (Jackson Immuno Research, Catalog No. 001-000-162) for 1 hour at room temperature and subsequently incubated with the respective primary antibody overnight at 4°C. The following primary antibodies were used: rabbit polyclonal ERF (Thermo Scientific, Catalog No. PA5-30237, 1:1,000), rabbit polyclonal SOX12 (Thermo Scientific, Catalog No. 23939-1-AP, 1:1,000), rabbit polyclonal CHCHD3 (Thermo Scientific, Catalog No. PA5-60167, 1:1,000), rabbit polyclonal PA2G4 (Thermo Scientific, Catalog No. 15348-1-AP, 1:1,000), and rabbit polyclonal TFAP4 (Thermo Scientific, Catalog No. 12017-1-AP, 1:1,000). Primary antibody was detected with 1 hour incubation at room temperature with HRP-linked goat anti–rabbit IgG (Cell Signaling Technology, Catalog No. 7074S,1:10,000) together with SuperSignal West Pico Plus Chemiluminescent Substrate (Thermo Scientific, Catalog No. 34578). For staining of β-Actin, blots were stripped for 15 minutes at room temperature using 1X ReBlot Plus Strong Antibody Stripping Solution (Millipore Sigma, Catalog No. 2504), re-blocked with 5% BSA (Jackson Immuno Research, Catalog No. 001-000-162) for 1 hour at room temperature and subsequently incubated with Direct-Blot HRP anti-β-Actin (BioLegend, Catalog No. 664803, 1:100,000) for 30 minutes at room temperature followed by SuperSignal West Pico Plus Chemiluminescent Substrate (Thermo Scientific, Catalog No. 34578). A ChemiDoc XRS+ (BioRad) was used to visualize protein bands, which were subsequently quantified with ImageJ software and normalized to the corresponding β-Actin bands.

### qPCR for senescence markers

RNA (200 ng) was reverse transcribed using the Applied Biosystems High-Capacity cDNA Reverse Transcription Kit (Thermo Scientific, Catalog No. 4368814) and the output volume of 20 μL was diluted in nuclease-free water to 40μL for a cDNA working concentration of 5ng/μL. Real time PCR was performed using the Applied Biosystems *Power* SYBR Green PCR Master Mix (Thermo Scientific, Catalog No. 4367659) on a Quant Studio 3 (Applied Biosystems) with reaction specificity confirmed by melt curve analysis. All comparisons (siSCR vs. master regulator knockdown) for each qPCR reaction were run on the same qPCR plate and run in a triplicate. Human qPCR primer sequences used (5’ to 3’) are listed in Table 1.

**Table 1.**
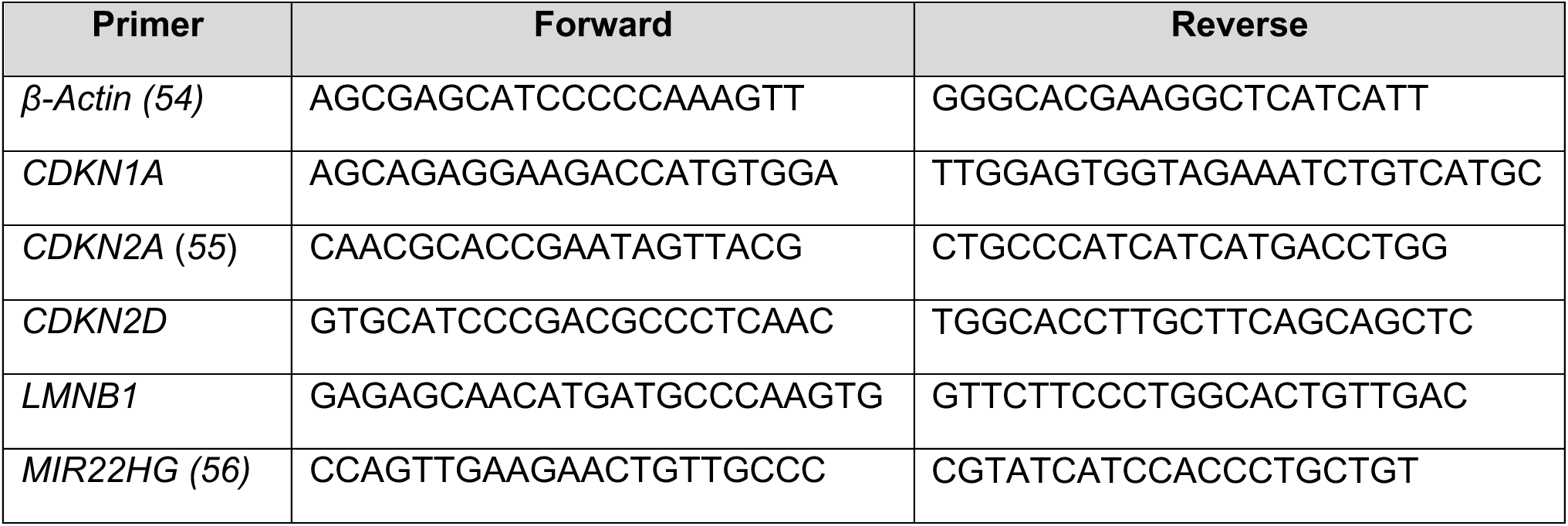
X

### Senolytic treatment in human cell lines

Human cell lines were plated on 96-well plates and treated for 7-days with DMSO or 100 uM BrdU as described above. Viability of the cells was then measured (see below) before senolytic drugs were added. Dasatinib (VWR, Catalog No. 102541-694) was dissolved in sterile DMSO to a stock concentration of 500 uM. The drug was then diluted to a working stock concentration of 2 uM in the appropriate culture media based on the cell type being tested before use in final concentrations of 50 nM or 100 nM. Quercetin (Med Chem Express, Catalog No. HY-18085) was dissolved in sterile DMSO to a stock concentration of 100 mM. The drug was then diluted to a working stock concentration of 2 mM in the appropriate culture media based on the cell type being tested before use in final concentrations of 10 uM or 20 uM. Navitoclax (ABT-263, Med Chem Express, Catalog No. HY-10087) was dissolved in sterile DMSO to a stock concentration of 20 mM. The drug was then diluted to a working stock concentration of 1 mM in the appropriate culture media based on the cell type being tested before use in final concentrations of 1 uM, 10 uM, or 20 uM. For all senolytic drugs used, an appropriate DMSO control (with the highest % DMSO used for each drug) was applied. All senolytics were applied for 48 hours and cells were incubated at 37°C with 5% CO_2_.

### Viability assay

To assess cellular viability, a CCK8 cell viability kit (Dojindo, Catalog No. CK04-05) was used according to the manufacturer’s protocol. In brief, cell medium was replaced by fresh medium containing 10% of CCK8 solution in the appropriate serum-containing media based on the cell type being tested. Using a SpectraMax iD3 (Molecular Devices) plate reader, an initial background absorbance reading at 450 nm was taken. Cells were then incubated for 2 hours at 37°C with 5% CO_2_ before absorbance was read again. The CCK8 assay for assessing viability was first used after the 7-day DMSO and BrdU treatment. Cells were then washed with PBS and senolytic drugs were added (as described above) and cells were incubated for 48 hours at 37°C with 5% CO_2_. The CCK8 assay was then performed again to assess viability based on treatment with senolytics and results were normalized to the initial CCK8 assay performed on Day 7 following DMSO or 100 uM BrdU treatment.

### Statistics

Statistical tests were performed using GraphPad’s Prism (v10) software. A threshold of p<0.05 was considered significant. Significance was determined using the test indicated in figure legends. Results are presented as means ± SD, with individual points representing biological replicates.

Representative growth curves in Figure S1A are associated with Figure 1C. Representative images in Figure S1B are associated with Figure 1D-F. A representative diagram in Figure S2A is associated with Figure 2B. Representative images in Figure S2B,C are associated with Figure 2D,E. Representative western blots of knockdowns in each cell type in Figure S6A-E are associated with knockdowns for Figure 6B-G and Figure 7C. Representative images in Figure S7 are associated with Figure 7F-J.

## Acknowledgements

This work was supported in part through grants 1R01NS124735 (MR), R01AG079898 (RBS), the Hartman Center (MR), the Center for Healthy Aging (MR), startup funds from Stony Brook University (RBS), and WaterWheel Foundation (RBS). Mitochondrial respiration measurements were performed using the Agilent Seahorse XFe96 analyzer with generous support from the Stony Brook University Biological Mass Spectrometry/Metabolomics core. Opinions, interpretations, conclusions, and recommendations are those of the author and are not necessarily endorsed by the sponsors.

## Author Contributions

Conceptualization: TR, MR; Methodology: TR, MR, JPB; Investigation: TR, MR, JPB; Visualization: TR, JPB; Funding acquisition: MR, RS; Project administration: MR, RS; Supervision: MR, RS; Writing – original draft: TR, JPB; Writing – review & editing: TR, MR, JPB, RS.

## Competing Interests

The authors declare no competing interests.

## SUPPLEMENTARY INFORMATION

**Figure S1:**
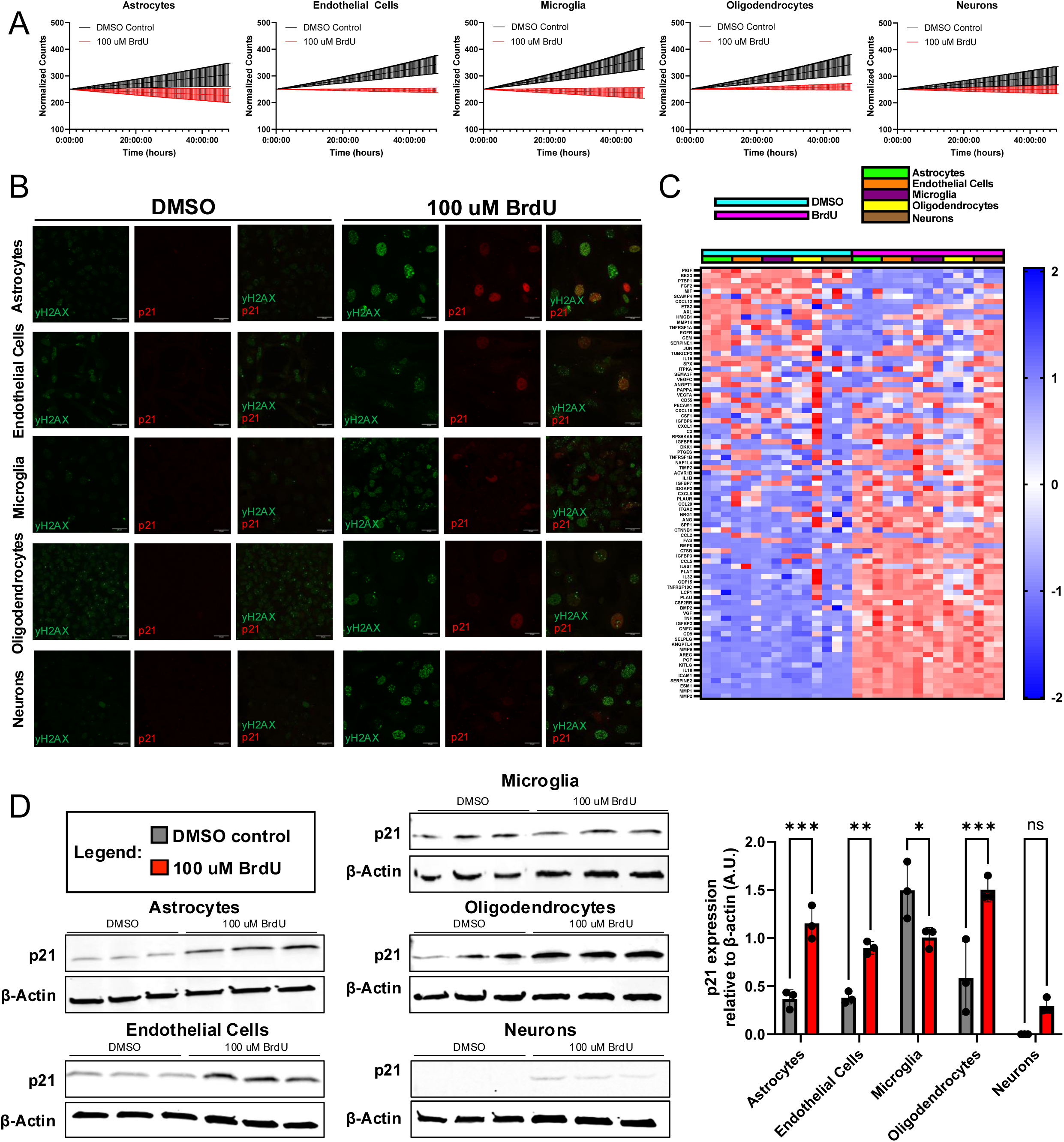
A) Normalized growth curves of DMSO (black) and BrdU (red) treated human cell lines across 48 hours. B) Confocal images of γH2AX (green) and p21 (red) staining after DMSO and 100 uM BrdU treatment in all cell lines. C) Z-score heatmap showing expression of established ‘SenMayo’ SASP factors in DMSO (blue) and BrdU (pink) treated human cell lines (astrocytes – green, endothelial cells – orange, microglia – purple, oligodendrocytes – yellow, neurons – brown). D) Western blots showing p21 expression levels in DMSO and 100 uM BrdU treated samples across all cell types and quantification of p21 levels relative to β-Actin levels from Western blots. (n=3). Data from (D) was analyzed by two-way ANOVA with Šídák’s multiple comparisons test. Bar graph in (D) shows mean with standard deviation (ns, p>0.05, * p<0.05, ** p<0.01). Scale bars are 33 um.

**Figure S2:**
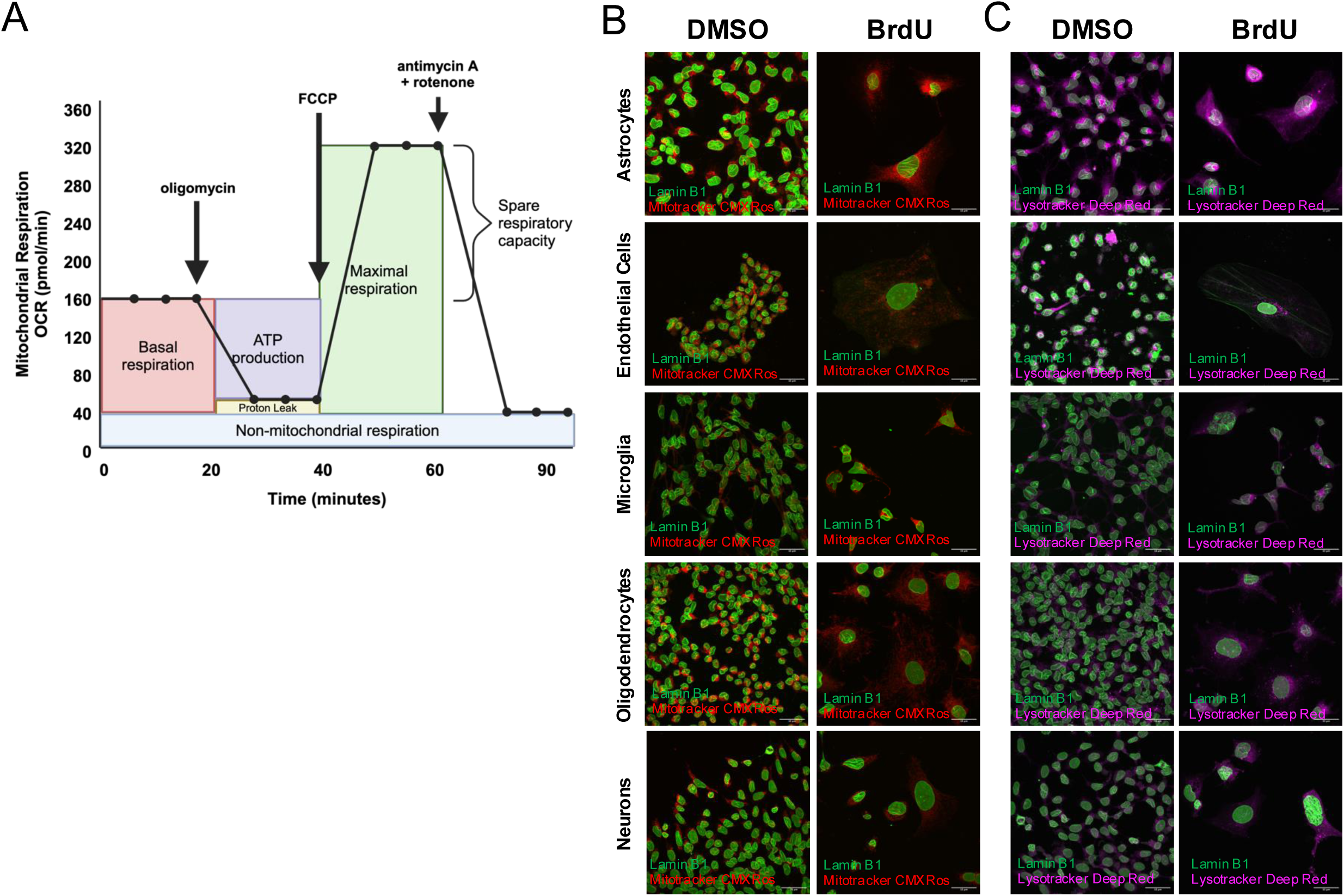
A) Overview of Agilent Seahorse XF Mito Stress Test protocol used for analyzing mitochondrial respiration of DMSO and BrdU treated cells (created with BioRender, Agilent). B) Confocal images of human cell lines (astrocytes, endothelial cells, microglia, oligodendrocytes, and neurons) treated with DMSO and 100 uM BrdU for 7-days with staining for lamin B1 (green) and Mitotracker Red CMXRos (red). C) Confocal images of human cell lines (astrocytes, endothelial cells, microglia, oligodendrocytes, and neurons) treated with DMSO and 100 uM BrdU for 7-days with staining for lamin B1 (green) and Lysotracker Deep Red (purple). Scale bars are 33 um.

**Figure S6:**
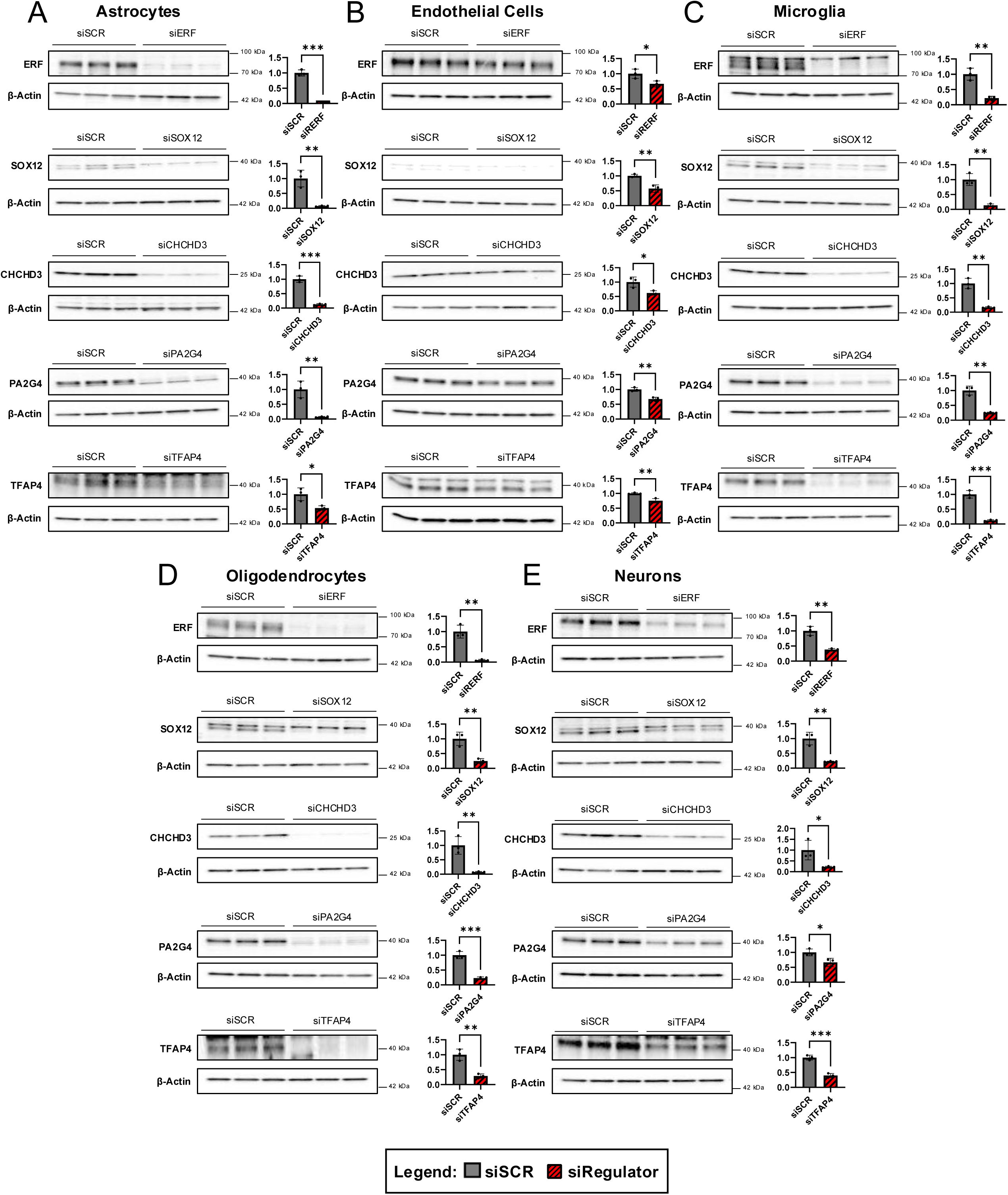
A-E) Representative Western blots (left) and respective quantification of normalized expression levels relative to β-Actin (right) showing knockdown of selected transcriptional regulators (red) (top-bottom): ERF, SOX12, CHCHD3, PA2G4, TFAP4 in astrocytes (A), endothelial cells (B), microglia (C), oligodendrocytes (D), and neurons (E) as compared to siSCR controls (grey). Data from (A-E) analyzed by unpaired two-tailed t-tests for each cell type and each transcriptional regulator (n=3).

**Figure S7:**
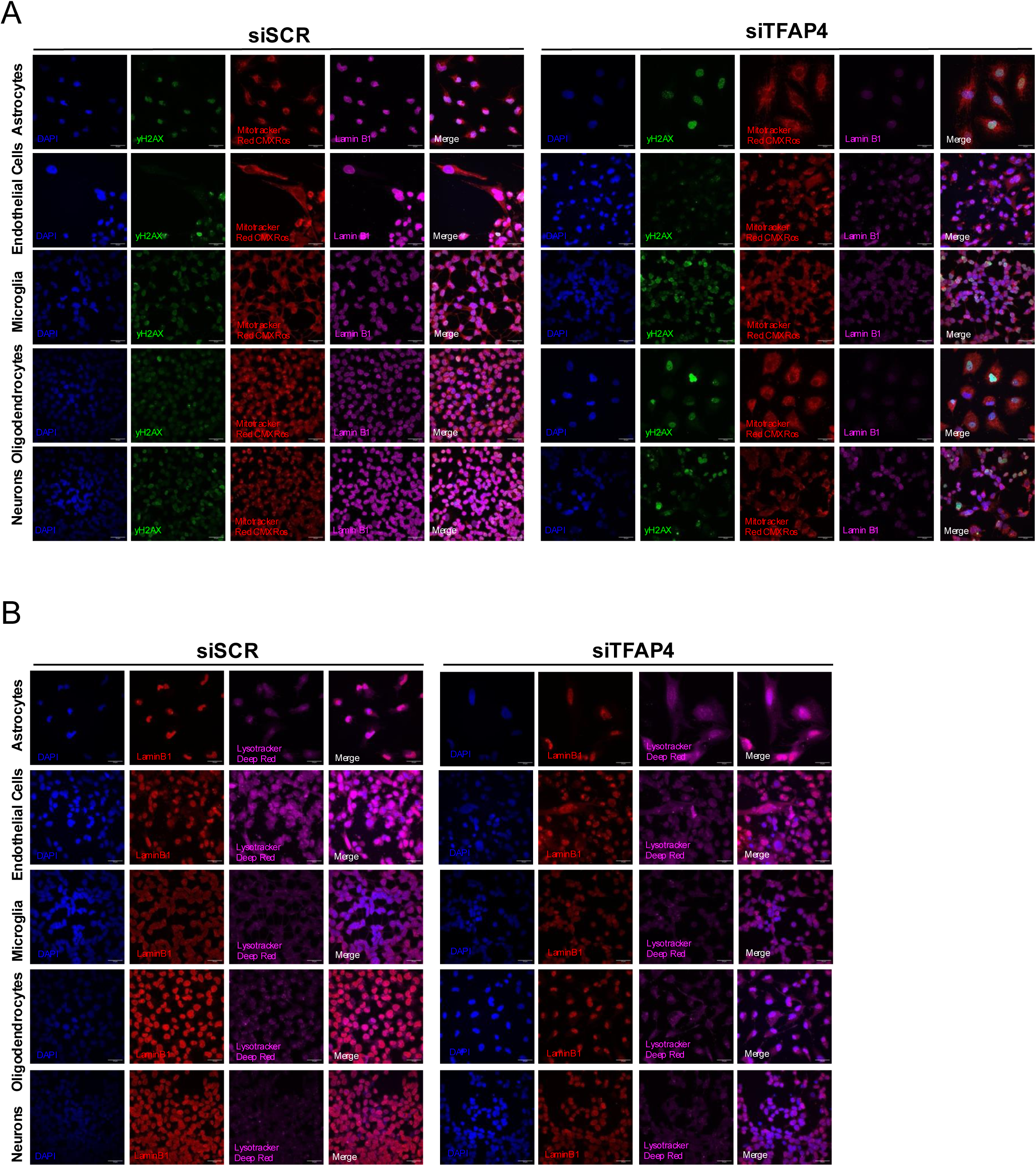
A) Confocal images of human cell lines treated with siSCR or siTFAP4 for with staining for γH2AX (green), Mitotracker Red CMXRos (red), and lamin B1 (purple) with DAPI (blue). B) Confocal images of human cell lines treated with siSCR or siTFAP4 with staining for lamin B1 (red) and Lysotracker Deep Red (purple) with DAPI (blue). Scale bars are 33 um.

## Notes

### Competing Interest Statement

The authors have declared no competing interest.

